# Telmisartan restricts Chikungunya virus infection *in vitro* and *in vivo* through the AT1/ PPAR-γ/MAPKs pathways

**DOI:** 10.1101/2021.07.30.454559

**Authors:** Saikat De, Prabhudutta Mamidi, Soumyajit Ghosh, Supriya Suman Keshry, Chandan Mahish, Sweta Smita Pani, Eshna Laha, Amrita Ray, Ankita Datey, Sanchari Chatterjee, Sharad Singh, Tathagata Mukherjee, Somlata Khamaru, Subhasis Chattopadhyay, Bharat Bhusan Subudhi, Soma Chattopadhyay

## Abstract

Chikungunya virus (CHIKV) has re-emerged as a global public health threat. The inflammatory pathways of RAS and PPAR-γ are usually involved in viral infections. Thus, Telmisartan (TM) with known capacity to block AT1 receptor and activate PPAR-γ, was investigated against CHIKV. The anti-CHIKV effect of TM was investigated *in vitro* (Vero, RAW 264.7 cells and hPBMCs) and *in vivo* (C57BL/6 mice). TM was found to abrogate CHIKV infection efficiently (IC50 of 15.34-20.89µM in the Vero and RAW 264.7 cells respectively). Viral RNA and proteins were reduced remarkably with the TM driven modulation of host m-TOR signaling. Additionally, TM interfered in the early and late stages of CHIKV life cycle with efficacy in both pre and post-treatment assay. Moreover, the agonist of AT1 receptor and antagonist of PPAR-γ increased CHIKV infection suggesting TM’s anti-viral potential by modulating host factors. Besides, reduced activation of all major MAPKs, NF-κB (p65) and cytokines by TM through the inflammatory axis supported the fact that the anti-CHIKV efficacy of TM is partly mediated through the AT1/PPAR-γ/MAPKs pathways. Interestingly, at the human equivalent dose, TM abrogated CHIKV infection and inflammation significantly leading to reduced clinical score and complete survival of C57BL/6 mice. Additionally, TM reduced infection in hPBMC derived monocyte-macrophage populations *in vitro*. Hence, TM was found to reduce CHIKV infection by targeting both viral and host factors. Considering its safety and *in vivo* efficacy, it can be a suitable candidate in future for repurposing against CHIKV.

## INTRODUCTION

Chikungunya virus (CHIKV) infection is categorized as a neglected tropical disease by WHO. However, in the last two decades, CHIKV has re-emerged and spread globally (1–4). CHIKV is an alphavirus transmitted to humans by the *Aedes sp*. of mosquitoes. The most common symptoms include fever, nausea, headaches, rash and poly-arthralgia. While the acute symptoms subside gradually, polyarthritis persists and may last for 90 days (5–8). Further, it is also known to cause neurological complications leading to irreversible brain damage (9). In the presence of co-morbidities and in vulnerable people, CHIKV infection can cause severe complications and death (10). Although mortality was underestimated earlier (11), between 2014-2017; more than 35000 deaths were associated to CHIKV infection across American and Caribbean regions (3). As no potent vaccine or drugs are available till date, extensive efforts are being taken to develop antiviral strategy for early clinical application (12, 13).

Drug repurposing is a strategy to find new clinical applications of the existing drugs with minimum cost, time and risk (14). Although drugs including chloroquine and ribavirin were repurposed against CHIKV, their therapeutic benefits are limited (13). Thus, there is a need for further investigation to find new drugs to reposition against CHIKV. Angiotensin II (Ang II) mediated activation of AT1 receptor in the Renin-Angiotensin System (RAS) is a primary mediator of oxidative stress and inflammation (15, 16). Its involvement in viral infection was first reported with improved survival of DBA/2 mice against Encephalomyocarditis (EMC) virus infection following administration of Ang-II inhibitors (17). Since then, it has been clinically correlated to viral diseases including Influenza (18), Bunya virus (19), Dengue virus (DENV) infections (20), Coxsackie virus (21) Ebola virus (22), Western Equine Encephalitis virus and neuro-adapted Sindbis virus (23). Considering the effects of Ang-II inhibitors against some of these alpha viruses, AT1 blocker drugs may be expected to be effective against CHIKV. Since CHIKV also affects CNS (24), it is desirable to use an AT1 blocker that can cross the blood-brain barrier. Only Telmisartan (TM) is known to have good brain permeability amongst this category of drugs (24). Our preliminary investigations suggested anti-CHIKV potential of TM (Indian Patent Application no. 201931012926, status: not published; filing Date: 30/03/2019). Recently, Tripathi et al. have corroborated this by demonstrating direct inhibition of the CHIKV-nsP2 protease activity as well as CHIKV infection in vitro (25). Thus, TM can be a potential drug for repurposing against CHIKV infection.

To justify the potential of TM for repurposing against CHIKV, the current investigation was carried out to understand its mode of action and effects *in vitro* and *in vivo* in mice. It was also investigated against other strains of CHIKV to justify its anti-CHIKV potential across different strains. The mode of action was investigated by looking at its impact on the levels of CHIKV RNA and proteins. Moreover, the interference in different phases of CHIKV life cycle was assessed. Finally, the effects were validated in the pre-clinical model in C57BL/6 mice using human equivalent doses and in hPBMC derived monocyte macrophage cells

## MATERIALS AND METHODS

### Virus and Cells

The Indian outbreak strain of CHIKV (IS, accession no. EF210157.2) and CHIKV prototype strain (PS, accession no. AF369024.2) were generous gifts from Dr. M.M. Parida (DRDE, Gwalior, India). Vero (African green monkey kidney epithelial cells) and RAW 264.7 (mouse monocyte/macrophage cells) cells were procured from NCCS, India. Vero cells were maintained in Dulbecco’s modified Eagle’s medium, (DMEM, PAN Biotech, Aiden Bach, Germany) supplemented with 10% Fetal bovine serum (FBS) (PAN Biotech, Aiden Bach, Germany), Gentamycin and Penicillin-streptomycin. RAW 264.7 cells were maintained in RPMI-1640 (Gibco RPMI-1640 GlutaMAX, Invitrogen, Cal, US) supplemented with 10% FBS (Gibco FBS, Invitrogen, Cal, US) and Penicillin-Streptomycin (PAN Biotech, Aidenbach, Germany). All the cells were maintained at 37°C with 5% CO_2_ in a humidified incubator.

### Antibodies and Inhibitors

The anti-CHIKV-E2 monoclonal antibody was a kind gift from Dr. M.M. Parida (DRDE, Gwalior, India). The mouse anti-CHIKV-nsP2 monoclonal antibody was developed by our group (26). AT1 monoclonal antibody was procured from Santa Cruz Biotech (Texas, US) and PPAR-γ polyclonal antibody was obtained from Cell Signalling Technology (CST) (MA, US). The antibodies for p38, p-p38, JNK, p-JNK, ERK1/2, p-ERK1/2, p-cJUN, p-IRF3, p-NF-κB, NF-κB, COX-2, p-mTOR, mTOR, p-4E-BP1, 4E-BP1, p-P70S6K and P70S6K were purchased from CST (MA, USA). The GAPDH and Actin antibodies were purchased from Abgenex India Pvt. Ltd. (Bhubaneswar, India) and Sigma Aldrich (Missouri, US) respectively. [Val5]-Angiotensin II acetate salt hydrate (Sigma Aldrich, Missouri, US) is defined as AG and GW9662 (Sigma Aldrich, Missouri,US) is abbreviated as GW throughout the study. Rosiglitazone (Rosi) and T0070907 (T007) were also procured from Sigma Aldrich. Telmisartan was a kind gift from Glenmark Life Sciences Ltd., Ankleshwar, Gujrat, India. Fluorochrome conjugated anti-human CD3, CD11b and CD19 antibodies were obtained from Abgenex, India Pvt. Ltd and anti-human CD14 antibody was purchased from eBioscience (San Diego, USA).

### Cytotoxicity Assay

MTT assay was performed to determine the cytotoxicity of TM, AG, GW, Rosi and T007 using EZcount™ MTT cell assay kit (Himedia, Mumbai, India) in Vero and RAW 264.7 cells according to the manufacturer’s protocol. Approximately 30000 numbers of cells were seeded in 96 well plates 24h before the experiment. After reaching 70% confluency, cells were treated with increasing concentrations of drugs/inhibitors along with DMSO as a reagent control. For cytotoxicity assay, 10μl MTT reagent (5mg/mL) was added to the cells 24h post drug treatment and incubated for 1h at 37ᵒC. Next, the media was removed and 100μl of solubilization buffer was added followed by incubation at 37°C for 15min to dissolve the Formazan crystals. Finally, the absorbance was measured at 570nm using a multimode plate reader and the metabolically active cell percentage was compared with the control cells to determine the cellular cytotoxicity (27).

### Virus infection

At 80% confluency, Vero cells were infected with PS at MOI 0.1 whereas, RAW 264.7 cells were infected with IS at MOI 5 followed by 90min incubation with shaking at every 10 to 15min interval. After infection, the inoculum was replaced with fresh media with different concentrations of drugs for 18h and 8h of PS infected Vero cells and IS infected RAW 264.7 cells respectively (27, 28). The cytopathic effect (CPE) was observed under microscope (20X magnification). Next, Infected/TM treated cells as well as supernatants were harvested for estimating the levels of viral RNA, proteins and viral titers according to the methods described earlier (29).

### Plaque Assay

Vero cells were seeded in 12-well plates and were infected as mentioned above. Mock, infected, and drug treated cell supernatants from different experiments were diluted and used for the infection of fresh Vero cells (70% confluent) seeded in 12 well plates. After 90mins, cells were washed with 1X PBS and overlaid with DMEM containing methylcellulose (Sigma, St. Louis, MO, USA) followed by incubation at 37ᵒC with 5% CO_2_ for 3 to 4 days. Then the cells were fixed with 8% formaldehyde, washed mildly with distilled water and stained with 0.1% crystal violet. The numbers of plaques were counted to calculate the virus titers which were presented as plaque-forming units per milliliter (pfu/mL) (27).

### Time of Addition Experiment

After infecting the cells with CHIKV (MOI 0.1), TM (100µM) was added to the cells at 0, 2, 4, 6, 8, 10, 12, 14 and 16hpi. Ribavirin (10μM) was used as a control. The supernatants were collected at 20hpi and viral titer was determined by plaque assay as mentioned before.

### Confocal Microscopy

Vero cells were seeded on coverslips in 6-well plates. After reaching 50% confluency, cells were infected as described earlier. At 16hpi (hour post infection), cells were washed thrice with 1X PBS gently, followed by fixation in 4% paraformaldehyde for 30min at RT and washed with 1X PBS. To permeabilize the cells, 0.5% Triton-X-100 was added to each well for 5min. After three washes, cells were blocked with 3% BSA for 30min at RT and incubated with primary antibody of viral protein E2 (1:750) for 1h. After primary antibody incubation, secondary anti-rabbit Alexa Flour 594 antibody was added at 1:1000 and 1:750 dilutions, respectively for 45min. The cells were stained with 4, 6-diamidino-2-phenylindole (DAPI; Life technology) and mounted with antifade reagent (Invitrogen). The fluorescence microscopic images were acquired using the Leica TCS SP5 confocal microscope (Leica Microsystems, Heidelberg, Germany) with 20X objective. The images were captured and analyzed using the Leica Application Suite Advanced Fluorescence (LASAF) V.1.8.1 software (30).

### Animal Studies

Animal experiments were conducted strictly under the guidelines defined by The Committee for the Purpose of Control and Supervision of Experiments on Animals (CPCSEA) of India. All procedures and experiments were reviewed and approved by the Institutional Animal Ethics Committee (76/Go/Rebi/S/1999/CPCSEA, 28.02.17). The 10-12 days old C57BL/6 mice were bred and housed under specific-pathogen-free conditions at our Animal House facility. For CHIKV infection, 10-12 days old mice were infected with 1×10^7^ pfu of CHIKV-PS subcutaneously at the flank region of the right hind limb. Serum free media was injected in the same region to the mock mice. Two hours post infection, 10mg/Kg TM was given orally to the treated group of mice (n=3). This was continued at every 12h interval up to 4 or 5 day post infection (dpi), whereas the mock and infection-control group (n=3) received only solvent. All the animals were monitored for disease symptoms every day. On 5 or 6dpi, depending upon the symptoms, mice were sacrificed and serum was isolated from the blood. Different tissues were also collected and stored in RNA later for RNA isolation, in 10% formalin for histological analysis and were snap frozen in liquid nitrogen for Western blot analysis. For the survival curve and clinical score studies the above mentioned infection and treatment protocols were followed (n=6 mice in three groups). However, drug was administrated up to 8dpi. For this study, all mice were monitored every day up to 8dpi. Clinical score of each mice were tabulated on daily basis according to the symptom based disease outcomes [no symptoms-0, fur rise-1, hunchback-2, one hind limb paralysis-3, both hind limb paralysis-4, death-5]. Mice mortality was also noted for the survival curve analysis (31).

### qRT-PCR

Vero cells were seeded and infected with CHIKV as described above. Viral RNA was extracted from the cells as well as supernatants of infected and treated samples. Cells were harvested and RNA extraction was performed using the Trizol reagent (Invitrogen, MA, US) (27). The cDNA synthesis was carried out with an equal amount of RNA by using the First Strand cDNA synthesis kit (Invitrogen, MA, US) and qRT-PCR was performed using specific primers for the CHIKV-E1 and nsP2 genes (Table. S1). GAPDH was taken as an endogenous control. On the other hand, equal volume of supernatant or serum samples were collected for viral RNA extraction using the QIAamp Viral RNA Mini kit (Qiagen, Hilden, Germany) according to the manufacturer’s instructions. An equal volume of RNA was used to synthesize cDNA and the E1 gene was amplified by qRT-PCR with equal volume of cDNA. The Ct values were plotted against the standard curve to obtain CHIKV copy number (29).

### Western blot

Protein expression was examined by the Western blot analysis. In brief, virus infected and drug treated Vero and RAW 264.7 cells were harvested at different hpi and lysed subsequently with equal volume of RIPA buffer. Snap frozen mice tissue samples were homogenized by hand homogenizer and lysed in RIPA buffer using syringe lysis method. Equal concentration of protein was separated on 10% SDS-polyacrylamide gel and was transferred onto PVDF membrane. Next the membrane was probed with anti-nsP2 (1:1000), anti-E2 (1:2500), anti-AT1 (1:250) monoclonal antibodies and anti-PPAR-γ (1:500) polyclonal antibody. All major MAPKs (p38 pAb, p-p38 pAb, JNK mAb, p-JNK mAb, ERK1/2 mAb, p-ERK1/2 mAb), p-cJUN mAb, p-IRF3 mAb, p-NF-κB mAb, NF-κB mAb, COX-2 mAb, p-mTOR mAb, mTOR mAb, p-4E-BP1 mAb, 4E-BP1 mAb, p-P70S6K pAb and P70S6K mAb were used in 1:1000 dilution to probe the membrane. GAPDH monoclonal (1:3000) and actin polyclonal (1:500) antibodies were used as loading control. Blots were developed by Immobilon Western Chemiluminiscent HRP substrate (Millipore, MA, US) in ChemiDoc MP Imaging System (Bio-Rad Laboratories, CAL, US) and intensities of all protein bands were quantified from three independent experiments using the Image J software (32).

### Flow cytometry

Flow cytometry was performed as previously described by Mishra et al (27). In brief, mock, CHIKV-infected and drug treated cells were harvested by cellular scrapping. Fixing the cells was performed with 4% paraformaldehyde for 10 min at RT followed by washing with 1X PBS. Then the cells were re-suspended in FACS buffer (1X PBS, 1%BSA, 0.01%NaN_3_) for staining. Permeabilization buffer (1XPBS + 0.5% BSA + 0.1% Saponin + 0.01% NaN_3_) was added to permeabilize the cells for intra cellular staining (ICS) of CHIKV antigen. Blocking of the cells was performed by 1% BSA for 30 min at RT. Primary antibodies (anti-mouse nsP2 and E2 mAbs) were then added for 1h followed by incubation with Alexa Fluor (AF) 488-conjugated chicken anti-mouse antibody (Invitrogen, USA). Approximately 1×10^4^ cells were acquired by the LSR-Fortessa flow cytometer (BD Biosciences, San Jose, California, USA) for each sample and analyzed by the FlowJo software (BD Biosciences, San Jose, California, USA)(27, 28).

### Sandwich ELISA

Cell culture supernatants were collected from different experimental sets and stored at −80°C. BD OptEIA™ Sandwich ELISA Kit (San Jose, CA, U.S.A.) for tumor necrosis factor alpha (TNFα) and interleukin-6 (IL-6) were used to determine cytokine levels in different test samples as per protocol described earlier(28, 32). Concentration of cytokines in different test samples were determined with respect to standard curves made from known concentrations of corresponding recombinant cytokines. Epoch 2 (Bio Tek, U.S.A.) microplate reader was used to determine the cytokine concentrations at 450 nm.

### Histopathological Examinations

For histopathological examinations, formalin-fixed tissue samples were dehydrated and embedded in paraffin wax, and serial paraffin sections of 5μM were obtained (33). The sections were then stained with hematoxylin-eosin (H&E), and histopathological changes were visualized using a light microscope (Zeiss Vert. A1, Germany). Sections were also examined for the presence of CHIKV-E2 protein using specific antibody. Briefly, E2 antibody was added to the slides for overnight at 4°C. After washing with PBS thrice, slides were incubated with Alexa Fluor 594 (anti mouse/ anti rabbit) for 45 minutes at room temperature in dark. The slides were washed with PBS thrice and then mounted with mounting reagent with DAPI and coverslips were applied to the slides (33).

### Measurement of mice cytokines levels by a Milliplex Map Kit

Serum samples were collected from CHIKV infected and drug treated 10-12 days old C57BL/6 mice at 5 or 6dpi. The Magnetic bead Milliplex Kit (Millipore, Billerica, MA, USA) was used to measure the levels of cytokines (IL-12(p40), MCP-1, IL-6, IP-10, KC, RANTES, GM-CSF, MIP-1β and TNF-α) according to the manufacturer’s instructions. The plate was read on Luminex-200TM with the xPONENT software (Luminex corporation, Austin, Texas, USA) (34).

### Isolation of human peripheral blood mononuclear cells (hPBMC)

For this 20mL blood was collected from three healthy donors in a 50mL centrifuge tube, rinsed with heparin and kept on ice till PBMC isolation. Isolated blood was diluted with an equal volume of chilled 1XPBS (HiMedia, India) followed by vigorous mixing. 2-3mL of Hi-Sep LSM (HiMedia, India) was poured in a 15mL tube and diluted blood was added in a drop-wise and slanted manner up to a final volume of 13mL per tube followed by centrifugation at 400RCF (25°C for 32min). Using Pasteur pipette, buffy coat was collected and cells were washed with 1XPBS twice at RT at 300RCF (25°C for 10min). Isolated PBMCs were plated in 6 well plates (Nunc, Thermo Fischer) at 5×10^6^ cells/well in RPMI 1640 (PAN biotech) supplemented with 15% FBS (PAN Biotech), Antibiotic-Antimycotic solution and L-Glutamine (HiMedia, India) for five days. Next, the cells were washed and supplemented with fresh culture medium in every two days interval (35, 36).

### Chikungunya virus infection in PBMC and flow cytometric staining

After five days, all adherent cells were detached using a cell scraper and seeded in a 24 well plate at a density of 0.15×10^6^. Cytotoxicity assay of TM was carried out for the adherent hPBMCs with various doses of TM by MTT assay as mentioned earlier and more than 95% cell viability was observed ranging up to 100µM concentration of TM. After one day, adherent cells were pre-treated with TM using 100µM concentration for 3h and subjected to CHIKV infection at 5 MOI for 2h. Infected cells were harvested at 12hpi followed by fixation with 4% PFA. Next, the cells were subjected to intracellular staining to detect viral protein E2 and surface staining for immunophenotyping of adherent population using flow cytometry. For immunophenotyping, fluorochrome conjugated anti-human CD3, CD11b, CD14 and CD19 antibodies were used (36, 37).

### Statistical analysis

All the statistical analyses were performed using the GraphPad Prism version 6.0.1 software. For mice experiment differences between mock, infected, and infected + drug treated groups were assessed by the one-way ANOVA Dunnett’s multiple comparisons test. P values are mean ± SE for n = 3 or 6.

## RESULTS

### TM inhibits CHIKV infection efficiently

In the MTT assay of TM (300µM), the viability of the Vero and the RAW 264.7 cells were found to be 100% and >90% respectively (Fig. 1A **&** 1F). Thus, a dose of 100µM or less was used in further investigations. To observe the effect of TM on reducing CPE, confluent Vero cells were infected with CHIKV PS at MOI 0.1. After 18hpi, a remarkable decrease in CPE was observed in cells treated with TM (30, 50, 70 and 100µM) as compared to the control (Fig. 1B). This anti-CHIKV potential was also evident from a subsequent study in confocal microscopy that showed a significant reduction in viral antigen E2 in a dose-dependent manner (Fig. 1C and D). The IC50 value of TM was found to be 15.34µM (Fig. 1E) and 20.89µM (Fig. 1G) against CHIKV-PS and CHIKV-IS respectively Hence, the data suggest the significant inhibitory capacity of TM to inhibit CHIKV infection *in vitro*.

**Figure 1.**
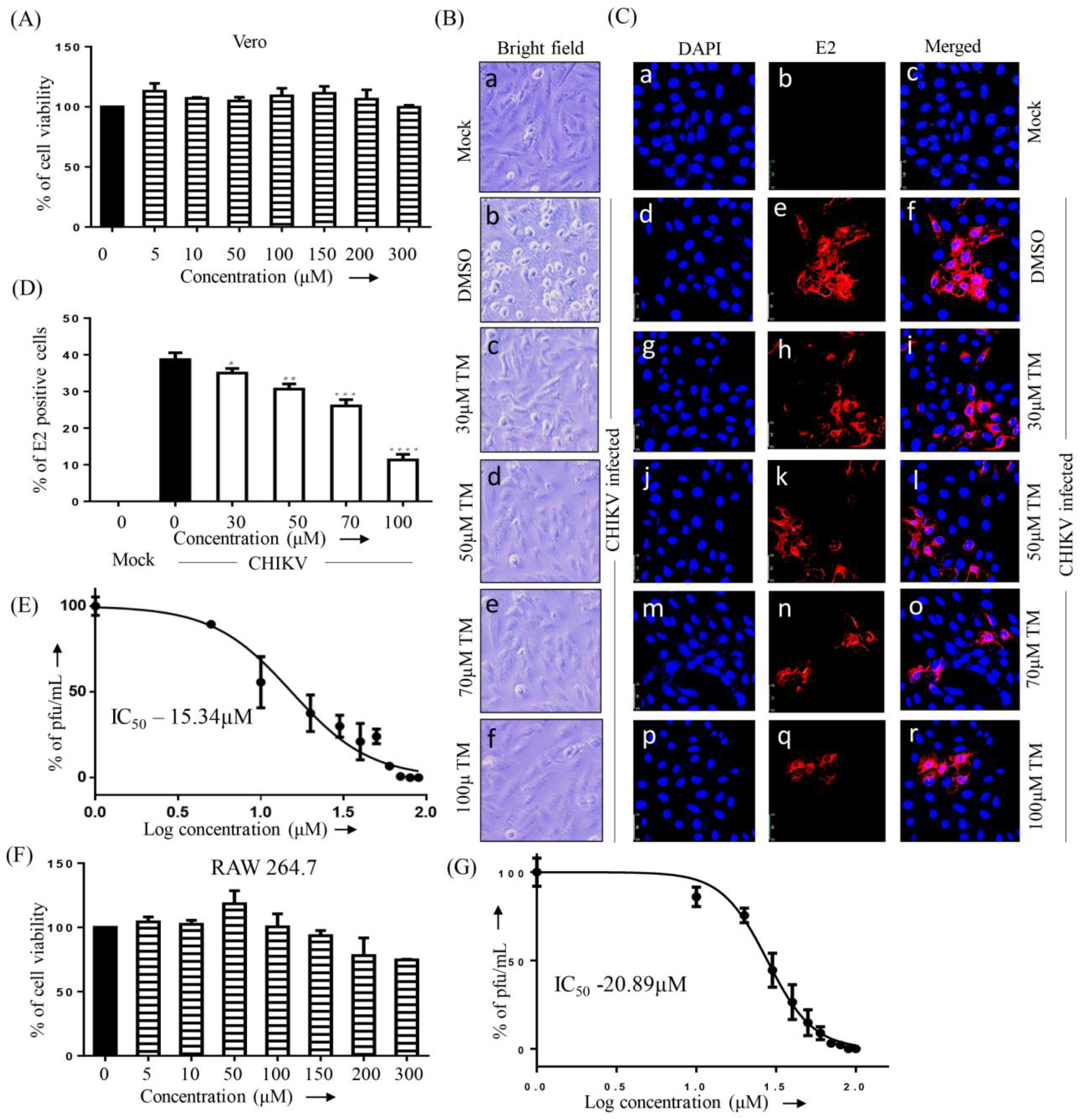
Telmisartan inhibits CHIKV infection efficiently. (A) Bar diagram showing the viability of Vero cells in presence of different concentration of TM. (B) Vero cells plated onto the cover-slips were infected with CHIKV-PS and TM was added with different concentrations (30µM, 50µM, 70µM, 100µM). CPE was observed under a microscope at 18hpi for both the uninfected and infected Vero cells after drug treatment and pictures were taken with 20X magnification in bright field microscope. (C) After 18hpi the cells were fixed and probed with E2 antibody followed by staining with secondary antibody, anti-mouse Alexa Fluor 594 (red) respectively. Nuclei were counterstained with DAPI (blue). (D) Bar diagram representing the percent positive E2 cell counts from the confocal images. (E) Vero cells were infected with CHIKV-PS and TM was added with different concentrations (10µM, 20µM, 30µM, 40µM, 50µM, 60µM, 70µM, 80µM, 90µM, 100µM). The supernatants were collected at 18hpi and virus titers were determined by plaque assay. The line diagram represents the IC50 value of TM in CHIKV-PS infected Vero cells where the X-axis depicts the logarithmic value of the different concentrations of TM and Y-axis depicts the percentage of pfu/mL. (F) Bar diagram showing the viability of RAW 264.7 cells in the presence of different concentration of TM. (G) RAW 264.7 cells were infected with CHIKV-IS and TM was added with different concentrations (10µM, 20µM, 30µM, 40µM, 50µM, 60µM, 70µM, 80µM, 90µM, 100µM). The supernatants were collected at 8hpi and virus titers were determined by plaque assay. The line diagram represents the IC50 value of TM in CHIKV-IS infected RAW 264.7 cells, where X and Y-axis represents the same as mentioned above. Data represented as mean ± SEM (n=3, p ≤ 0.05 was considered statistically significant)

### TM reduces the viral RNA and protein levels

To confirm the effect of TM on CHIKV structural (E1/E2) and non-structural (nsP2) RNAs, qRT-PCR was conducted with respective primers (Table. S1), using virus infected and TM treated cells. At 100µM concentration, it reduced the CHIKV-E1 and CHIKV-nsP2 RNA levels by 25 fold and 5 fold respectively in CHIKV-PS infected Vero cells (Fig. S1A). Similar reduction of 90 and 45-fold was observed in CHIKV-IS infected RAW 264.7 cells in presence of 70µM TM (Fig. S1B). Further, viral protein levels were estimated by Western blots and flow cytometry. At 100µM concentration, it reduced the nsP2 and E2 protein levels by 86% and 95% respectively in CHIKV-PS infected Vero cells (Fig. 2A, B, C **and** D). In CHIKV-IS infected RAW 264.7 cells, TM (70µM) reduced the nsP2 and E2 protein levels by 75% and 88% respectively (Fig. 2E, F, G **and** H). Thus, it could be suggested that TM inhibits CHIKV infection by reducing viral RNAs as well as proteins.

**Figure 2.**
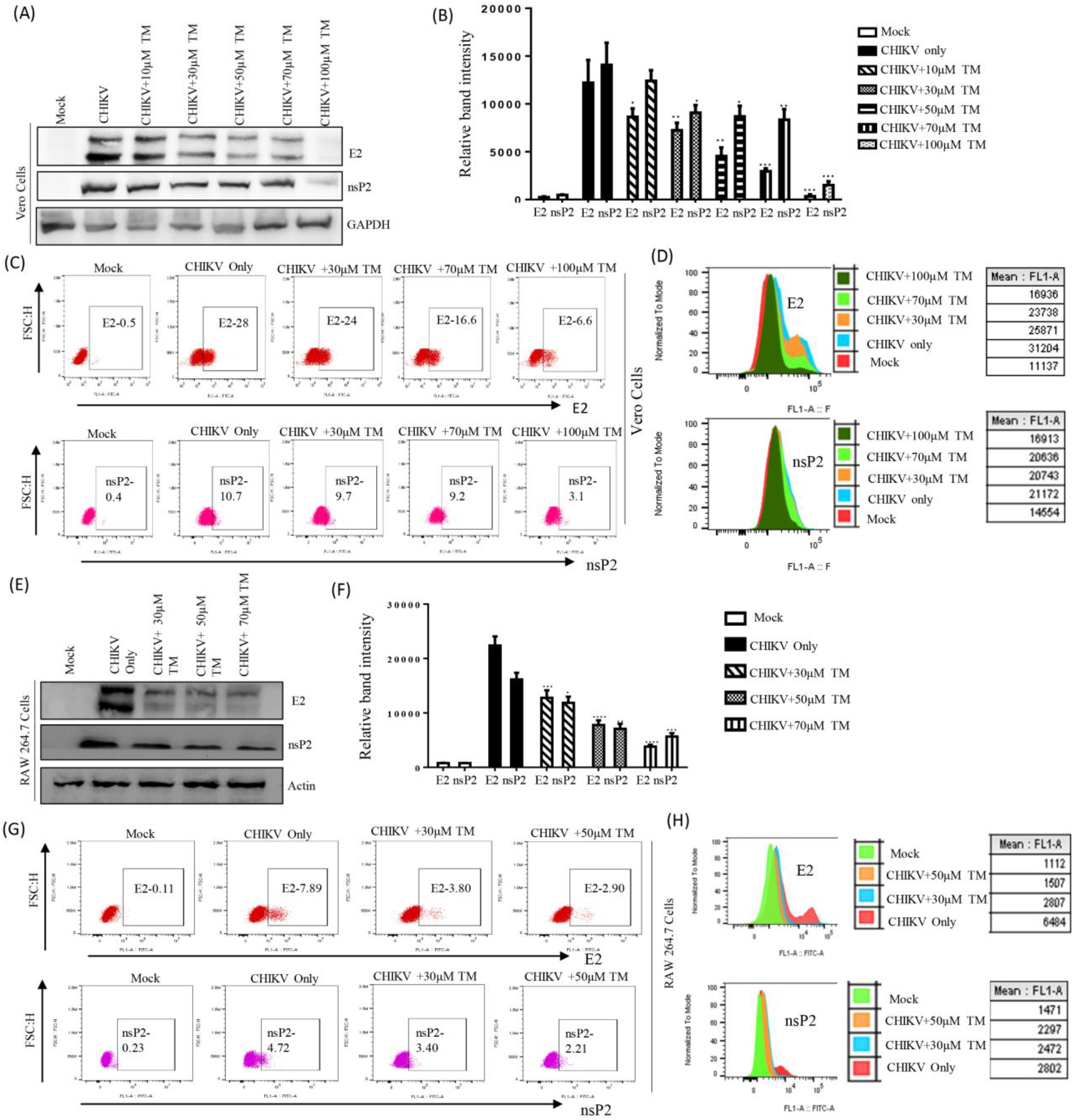
Reduction of CHIKV protein after TM treatment: Vero cells and RAW 264.7 cells were infected with CHIKV and treated with different concentrations of TM as mentioned above. Vero Cells were harvested at 18hpi, whereas RAW 264.7 cells were harvested at 8hpi. (A, E) The Vero and RAW 264.7 cell lysates were processed for Western blot using antibodies against CHIKV nsP2 and E2 proteins. GAPDH and Actin was used as a loading control for Vero cells and RAW 264.7 cells respectively. (B, F) Bar diagrams showing the relative band intensities of CHIKV-E2 and CHIKV-nsP2 protein expressions in Vero and RAW 264.7 cells respectively. Data represented as mean ± SEM (n=3, p ≤ 0.05 was considered statistically significant). (C, G) Cells were stained with CHIKV nsP2 and E2 proteins and analyzed in Flow Cytometry during infection in Vero cells and RAW 264.7 cells respectively. Dot plot analysis showing percent positive cells for E2 and nsP2 against FSC-H. (D, H) Histogram and table showing Mean fluorescent intensities (MFI) of E2 and nsP2 proteins.

### TM interferes in the early and late stages of CHIKV life cycle

To understand which stage of CHIKV life cycle is affected by TM, the “time-of-addition” experiment was performed. The estimated viral titers showed that the release of infectious virus particles was abrogated by 95-82% with the addition of TM at 0 to 10hpi as compared to control (Fig. 3A). Even after addition of TM at 12 to 16hpi, the abrogation of infectious virus particle release was 75-45%. The positive control (Ribavirin) is well known for interfering in the early phase of the CHIKV life cycle and accordingly, it exhibited 88-70% decrease in the release of infectious virus particles when it was added at 0 to 6hpi. However, very less reduction (10-0%) was observed at 12 to 16hpi compared to the infection control (Fig. 3A). Therefore, it can be concluded that TM might interfere in both the early and late stages of the CHIKV life cycle.

**Figure 3.**
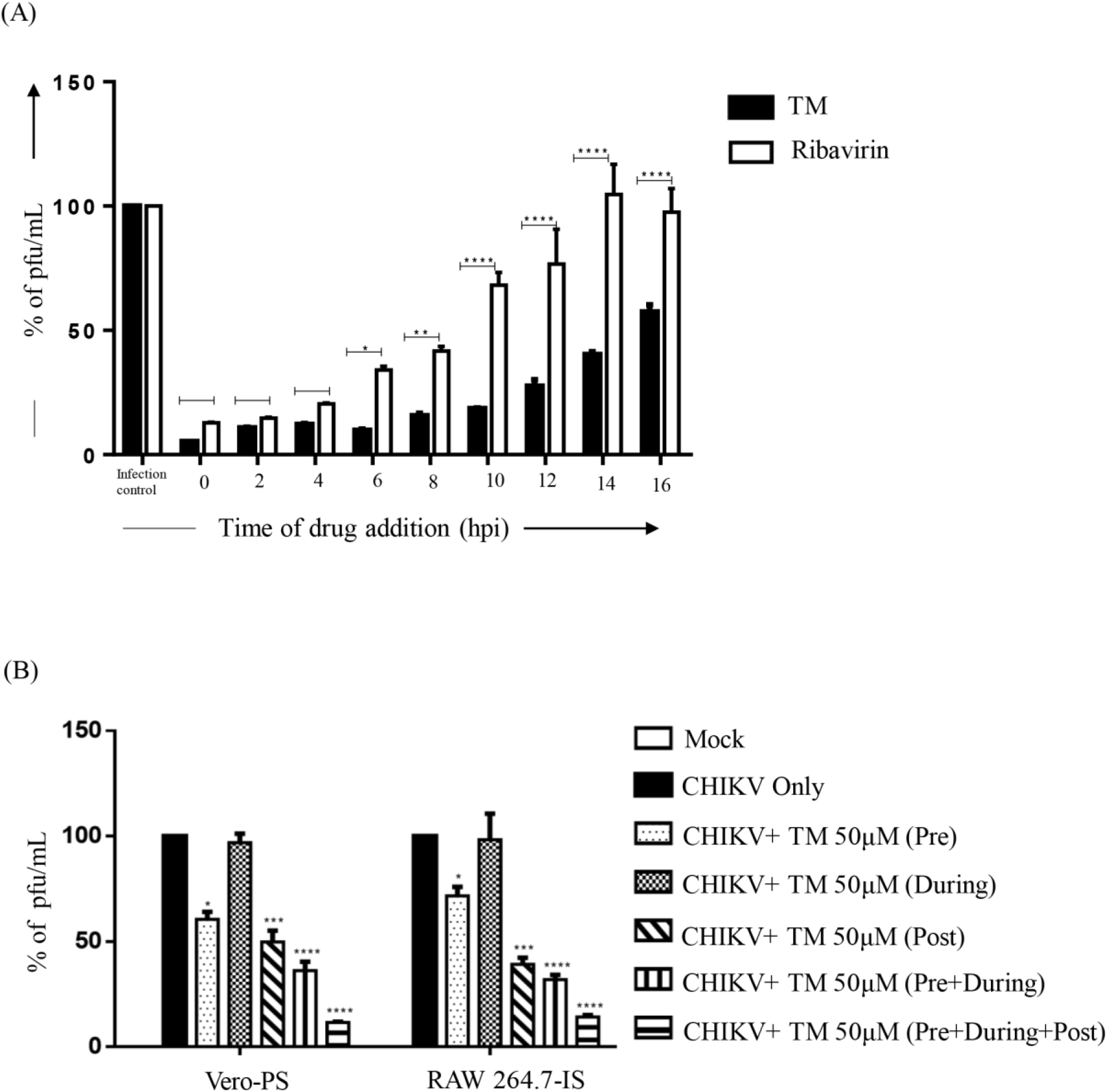
TM interferes in early and late stages of CHIKV life cycle. Pre and Post-treatments of TM also reduce CHIKV infection significantly. (A) Vero cells were infected by CHIKV-PS with MOI 0.1 and 100μM TM was added to each sample, every 2h interval up to 16hpi. Ribavirin (10μM) was used as a control. The bar diagram represents the percentage of virus titer of the supernatants collected at 18hpi for all the samples. (B) Vero and RAW 264.7 cells were treated with TM (50µM) separately before, during and post infection. Supernatants collected at 18 hpi for Vero and 8hpi for RAW 264.7 cells were subjected to plaque assay. Bar diagram showing percentage of virus particle. Data presented as mean ± SEM. (n=3, p ≤ 0.05 was considered statistically significant).

### Pre and post-treatment of TM reduces CHIKV infection significantly

To evaluate the effectiveness of various treatment regimens, TM (50μM) was administered at different stages of infection. The results reveal that viral particle formation was abrogated by 30 - 40% following pre-treatment with TM (Fig. 3B). Whereas, very little (3-4%) or almost no inhibition was found when TM was used during infection. Compared to these, TM was relatively more effective in the post-infection stage with a reduction of 55-64% in virus particle formation. Interestingly, 65-70% reduction was observed when the cells were treated with TM before and during the infection. However, it was most effective when added in pre, during and post infection with 87-90% abrogation of viral particle formation. The data suggest that TM might not have an effect on the attachment process; nevertheless, presence of TM in all the phases demonstrated the best effectiveness against CHIKV.

### Inhibition of CHIKV by TM is mediated through AT1 and PPAR-γ pathway

Since TM is a modulator of AT1 and PPAR-γ (20, 38), these host factors might have roles in its anti-CHIKV property. Thus, agonist of AT1 (AG) and antagonist of PPAR-γ (GW) were used in this study. GW and AG were non-toxic at concentration of 20μM (Fig. S2A **and** B) and 100μM (Fig. S2C **and** D) respectively. Their treatment before and after viral infection revealed that both GW (Fig. S2E **and** F) and AG (Fig. S2G **and** H) augmented viral particle formation dose dependently with up regulation of the CHIKV-E2 and nsP2 protein levels (Fig. S2I **and** J). Subsequently, this increased CPE and level of E2 protein (Fig. 4A). This was associated with enhanced CHIKV progeny release in presence of both GW [25% in Vero and 35% in RAW 264.7 cells; (Fig. 4B and C)] and AG [42% in Vero and 105% in RAW 264.7 cells; (Fig. 4B and C)]. Although significant increase in levels of CHIKV E2 and nsP2 was observed with both GW (Fig. 4D, E, H **and** I) and AG, the levels were relatively higher following AG treatment (Fig. 4D, E, H **and** I). In contrast, TM treatment significantly reduced infection as well as levels of CHIKV proteins. While AT1 protein level was up-regulated by 15-20% upon infection, TM treatment down-regulated this by 53-74% (Fig. 4D, F, H **and** J). On the other hand, there was modest down-regulation of PPAR-γ in infected cells and TM treatment augmented the PPAR-γ expression by 140-300% (Fig. 4D, G, H **and** K). To strengthen this observation, an established PPAR-γ agonist Rosiglitazone (Rosi) was tested against CHIKV. At a non-toxic dose of 200μM (Fig. S3A) it showed 93% inhibition of viral particle formation (Fig. S3C) with 60% up regulation of PPAR-γ along with reduction in the CHIKV-E2 (86%) and CHIKV-nsP2 (80%) protein levels (Fig. S3D **and** E). Further, a PPAR-γ inhibitor, T0070907 (T007) at a non-toxic concentration (Fig. S3B) of 20μM increased CHIKV particle formation by 35% (Fig. S3F) with up regulation of the CHIKV-E2 (62%) and CHIKV-nsP2 (51%) protein levels (Fig. S3G **and** H). Taken together, it could be suggested that the activation of the AT1 receptor and blocking of PPAR-γ have contributed to the enhancement of viral infection. Thus, the anti-CHIKV effect of TM can be mediated through these host factors.

**Figure 4.**
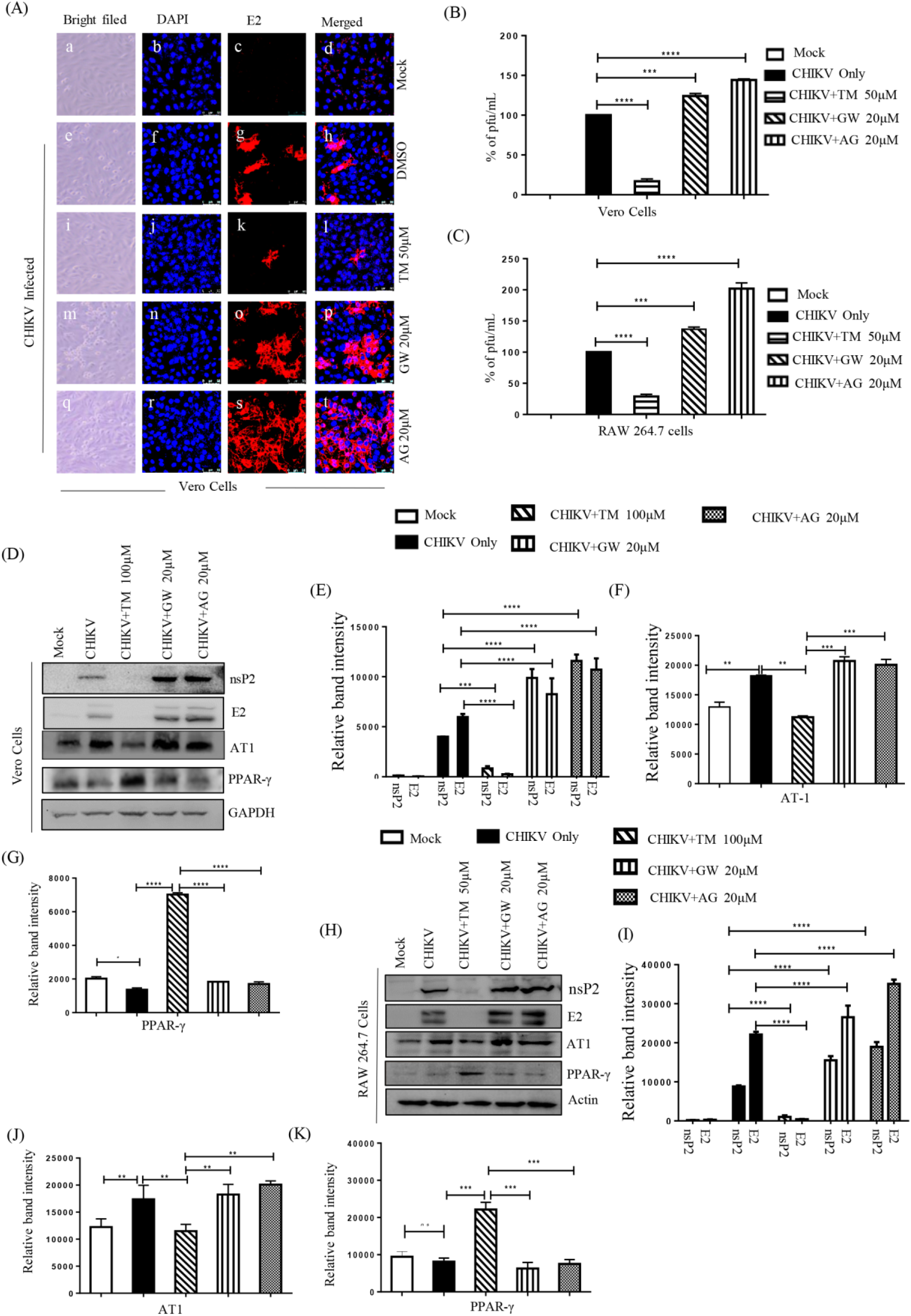
Inhibition of CHIKV by TM is mediated through AT1 and PPAR-γ. (A) Vero cells plated onto the cover-slips were treated with TM (50µM), GW (20µM), and AG (20µM) individually before the infection for 3 hours and infected with CHIKV-PS at MOI 0.1 and treated again with TM (50µM), GW (20µM), and AG (20µM) individually. At 18hpi, the cells were fixed and probed with E2 antibody followed by staining with secondary antibody, anti-mouse Alexa Fluor 594 (red). Nuclei were counterstained with DAPI (blue). (B) Bar diagrams depicting the percentage of viral titer of the collected supernatant. (C) RAW 264.7 cells were treated with TM (50µM), GW (20µM), AG (20µM) infected by CHIKV-IS at MOI-5 and again treated with TM (50µM), GW (20µM), AG (20µM). Bar diagrams indicating the percentage of viral titer in the cell supernatant. (D, H) Western blot showing the levels of different proteins (nsP2, E2, AT1 and PPAR-γ) in the infected cells. GAPDH and Actin was used as loading controls for Vero and RAW 264.7 cells respectively. Bar diagram indicating the relative band intensities of viral (nsp2 and E2) and host (AT1 and PPAR-γ) protein levels in Vero (E, F, G) and RAW 264.7 cells (I, J, K). Data presented as mean ± SEM. (n=3, p ≤ 0.05 was considered statistically significant).

### TM reduces CHIKV protein levels by modulating the host translational control pathway

To understand the mechanism behind the reduction in CHIKV protein levels by TM, virus infected RAW 264.7 cell lysates were analyzed for the expressions of host translational control pathway proteins. It was observed that the expression of p-mTOR, a kinase which regulates protein synthesis pathway (39) was reduced by 50% during TM treatment (Fig 5A and B). About 65% reduction was also observed in the expression of p-P70S6 kinase (Fig. 5A and D) that binds to ribosomal S6 protein to induces protein synthesis (40). Moreover, the expression of p4E-BP1 which binds to eukaryotic translational initiation factor 4E (EIF4E) and represses translation (41) was up regulated by around 45% (Fig. 5A and C). Hence, it might be speculated that TM reduces CHIKV protein levels by modulating the host translational control pathway.

**Figure 5.**
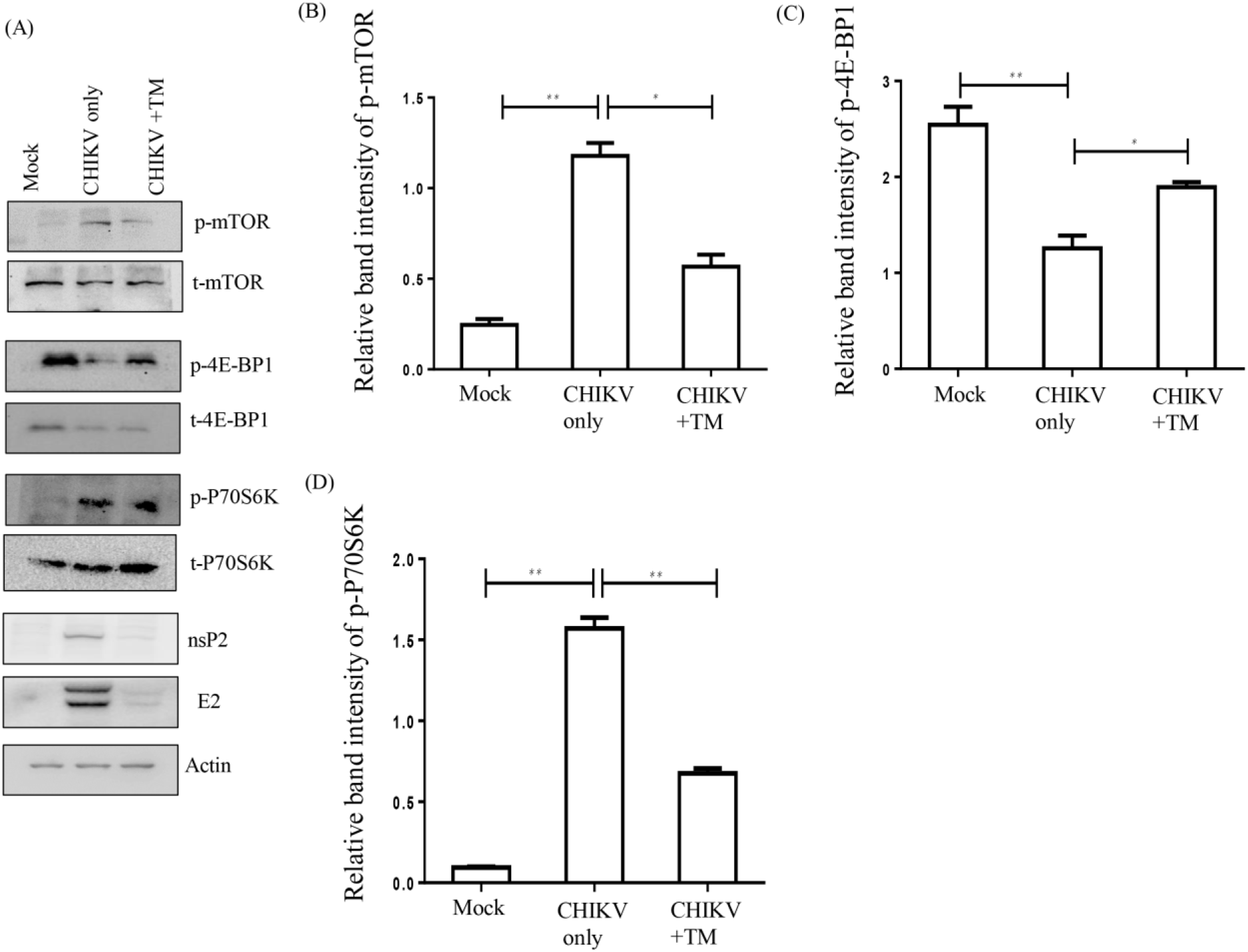
TM reduces CHIKV protein expression by modulating the host translational pathway. RAW 264.7 cells were CHIKV-IS infected, treated with TM (50μM), harvested at 8hpi (A) Western blot showing the phosphorylation status of mTOR, 4E-BP1, P70S6K and along with the viral protein (E2, nsP2) level in RAW 264.7 cell lysates. Actin was used as a loading control. (B, C, D) Bar diagrams indicating relative band intensities of p-mTOR, p-4E-BP1 and p-P70S6K respectively. Data presented as mean ± SEM. (n=3, p ≤ 0.05 was considered statistically significant).

### AT1 agonist (AG) and PPAR-γ antagonist (GW) mediated augmentation of CHIKV infection is also inhibited by TM

In order to understand the ability of TM to abrogate CHIKV infection that was augmented by AG or GW, RAW 264.7 cells were treated with AG or GW along with TM before and after CHIKV-IS infection. Interestingly, TM reduced infection by 80% and 70% when co-treated with GW or AG respectively (Fig. 6A). It also reduced the nsP2 protein level [85% in the presence of GW and 75% in the presence of AG; (Fig. 6B and C)]. The increase in the level of AT1 in presence of AG and decrease in the level of PPAR-γ in presence of GW was antagonized by TM with reduction in AT1 and enhancement in PPAR-γ levels (Fig. 6D and E). These results further support the fact that the anti-viral activity of TM is partly mediated through AT1 and PPAR-γ.

**Figure 6.**
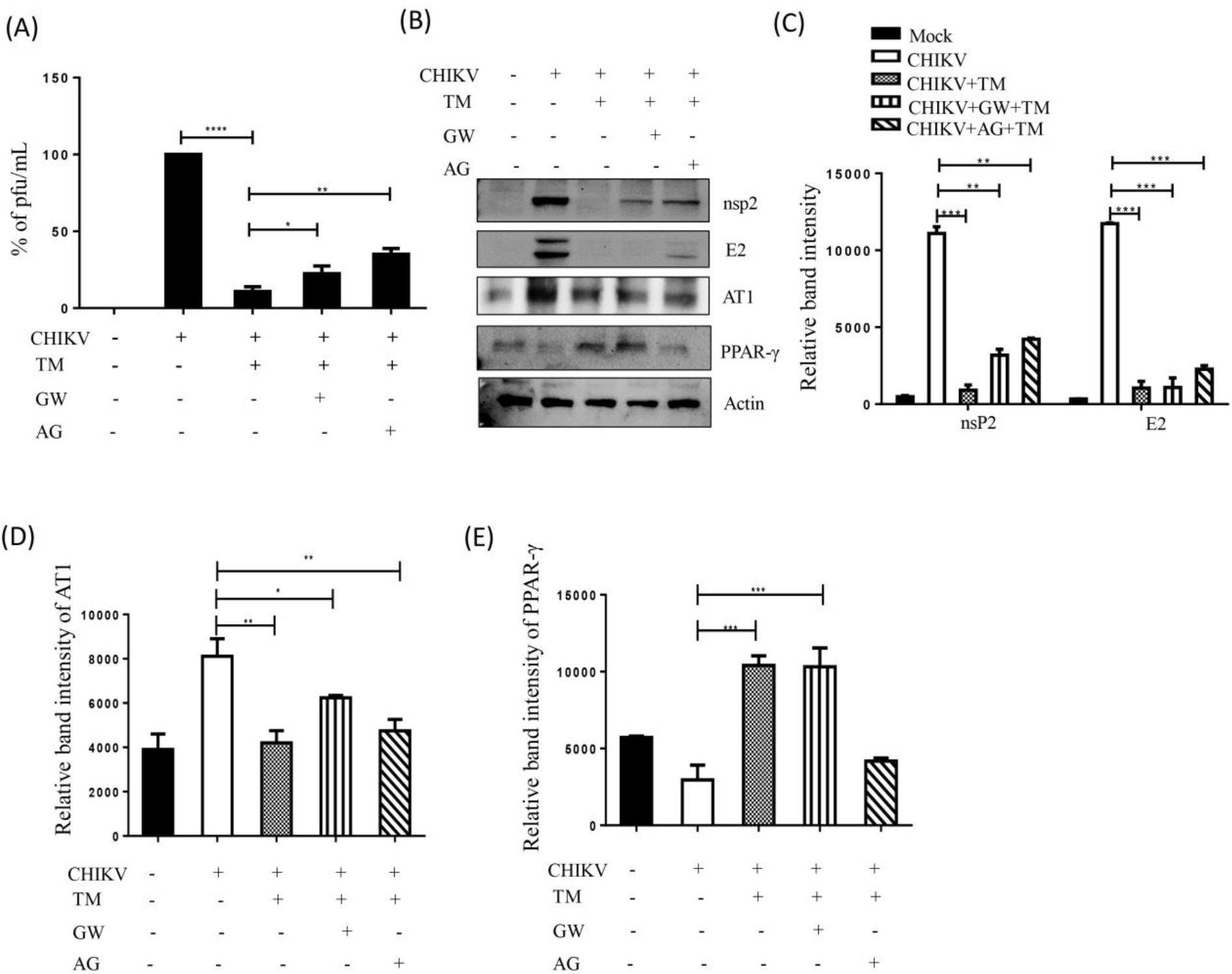
TM abrogates AT1 agonist (AG) and PPAR-γ antagonist (GW) mediated enhanced CHIKV infection. RAW 264.7 cells were co-treated with TM (50µM), GW (20µM), and AG (20µM) before and after CHIKV-IS infection at MOI-5. (A) Bar diagram indicating the percentage of viral titer in the cell supernatant. (B) Western blot showing the levels of different proteins (nsP2, E2, AT1 and PPAR-γ) in the infected cells. Actin was used as a loading control. Bar diagram indicating the relative band intensities of viral (nsp2 and E2) and host protein levels (AT1 and PPAR-γ) (C, D and E). Data presented as mean ± SEM. (n=3, p ≤ 0.05 was considered statistically significant).

### TM reduces the CHIKV induced inflammatory response through the MAPK pathway, NF-κB and COX-2

In order to find out whether TM modulates CHIKV induced inflammation, induction of p38, ERK1/2, JNK, cJUN, IRF3, Nf-κB, COX-2, TNF-α and IL-6 were estimated. Compared to mock, the CHIKV infection led to 2 to 3-fold induction in the phosphorylation of each of the major MAPKs: p38, ERK1/2, and JNK. However, treatment with TM resulted in reduced phosphorylation. Whereas, MAPKs were activated by 2.5 to 4-fold in infected cells compared to mock in presence of AG and GW (Fig. 7A, B, C **and** D). Phosphorylation of cJUN is one of the major transcription factors for the expression of proinflammatory cytokines (32). It’s induction was 10 fold higher after infection. While TM treatment reduced this level of cJUN by 2 fold, whereas, GW and AG increased it by 1.5 and 1.8 fold respectively (Fig. 7A and E). Furthermore, TM treatment reduced the expressions of another key transcription factor, p-IRF3 by 2 fold as compared to the infection control. Whereas, this level was observed to be 1.6 and 1.7 fold higher for GW and AG treatment respectively as compared to infection (Fig. 7A and F). Similarly, induction of p-NF-κB (p65) was down regulated by 2 fold in presence of TM but up regulated with GW and AG compared to infection (Fig. 7A and G). Also, expression of COX-2, a key enzyme that mediates prostaglandin synthesis (42), was down regulated by TM (2 fold), however, was up-regulated by GW and AG as compared to infection (Fig. 7H and I). TM treatment was also able to inhibit levels of cytokines significantly [TNF-α by 2.9 fold and IL-6 by 10 fold; (Fig. 7J and K)]. On the other hand, AG and GW treatment increased the levels of TNF-α and IL6 (data not shown). Altogether, reduction in activation of all major MAPKs and cytokines by TM through MAPKs inflammatory axis supports the fact that it’s anti-CHIKV property is partly mediated through the AT1/PPAR-γ/MAPKs pathways. Furthermore, down regulation of p-NF-κB and COX-2 indicates the anti-inflammatory effect of TM.

**Figure 7.**
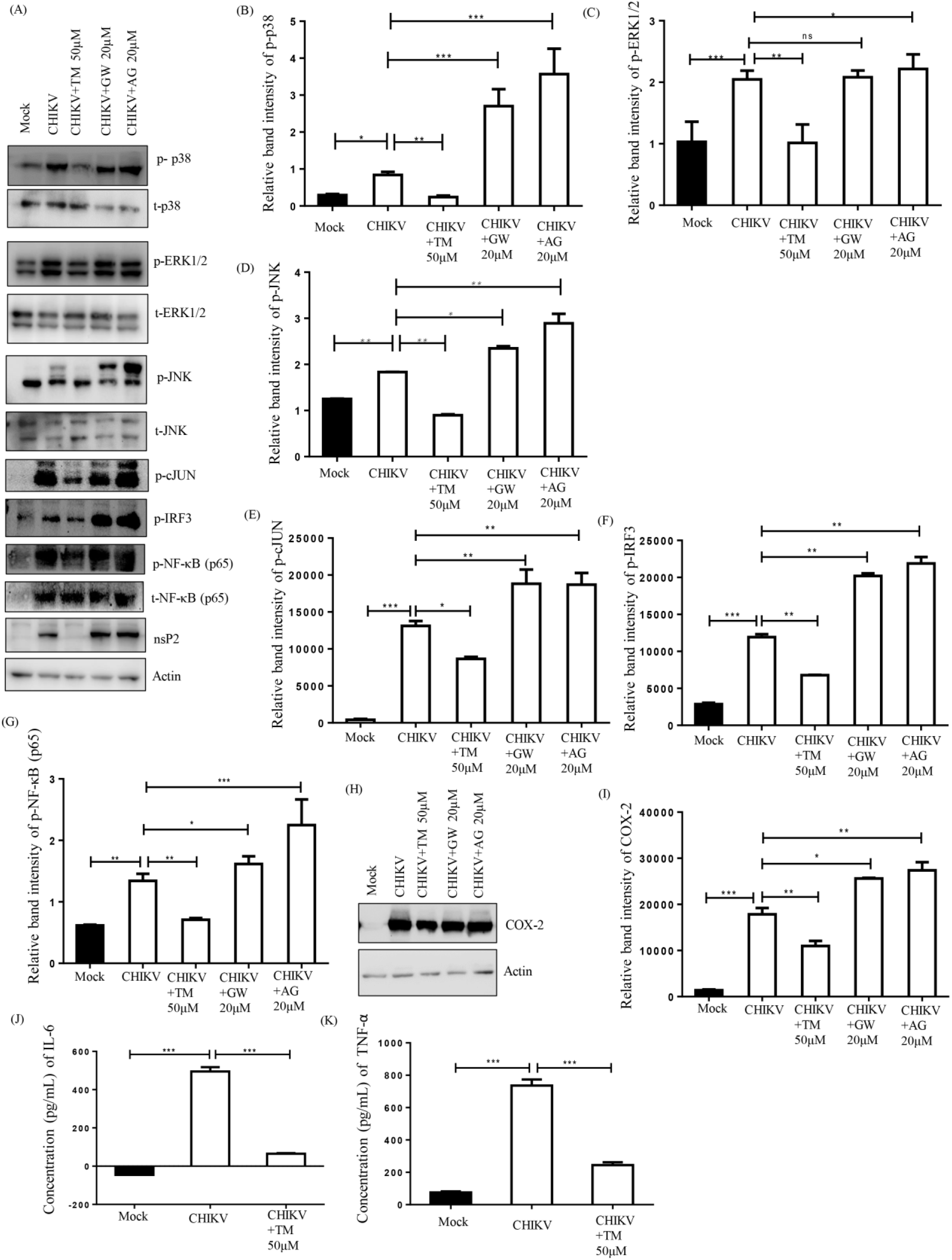
TM reduces the CHIKV induced inflammatory response through the MAPK pathway, NF-κB and COX-2. RAW 264.7 cells were infected with CHIKV-IS, treated with TM (50μM), GW (20μM) and AG (20μM) and were harvested at 8hpi. (A) Western blot showing the viral protein (nsP2) level along with phosphorylation status of p38, ERK, JNK, cJUN, IRF3 and NF-κB in RAW 264.7 cell lysates. Actin was used as a loading control. (B, C, D, E, F, G) Bar diagrams indicating relative band intensities of p-p38, p-ERK, p-JNK, p-cJUN, p-IRF3 and p-NF-κB. (H) Western blot showing the level of COX-2 protein. (I) Bar diagrams depicting relative band intensities of COX-2 protein. (J, K) Bar diagram showing the levels of secreted cytokines (TNF-α and IL-6) of CHIKV infected and TM treated macrophages in the supernatants quantified using sandwich ELISA of CHIKV infected and TM treated macrophages. Data represented mean ± SEM (n=3). p ≤ 0.05 was considered as statistically significant.

### TM protects mice from CHIKV infection and inflammation

The anti-CHIKV effect of TM was assessed in C57BL/6 mice infected with CHIKV-PS following treatment with TM (10mg/Kg) at every 12h interval up to 5 or 6dpi. The infected mice showed symptoms of arthritis in limbs and impaired mobility, whereas TM-treated animals showed no such abnormalities (Fig. 8A**, Movie. S1**). Based on viral RNA isolated from the serum of animals, the CHIKV copy number was reduced by 55% with TM (Fig. 8B). In muscles, reduction of in CHIKV-E1 RNA (4-fold) and in CHIKV-E2 protein (62%) was observed (Fig. 8C, D **and** E). Similarly, reduction of E1 RNA was detected in other organs like liver (5 fold), kidney (10 fold) and spleen (100 fold). The protein levels of CHIKV-E2 were also reduced in liver (40%), kidney (66%) and spleen (75%). Further, H&E stained infected muscle sections of mice showed huge infiltration of the mononuclear lymphocytes (Fig. 8F a-II) which was remarkably lower after TM treatment (Fig. 8F a-III). Additionally, IHC analysis revealed the reduction of E2 level in CHIKV infected muscle upon TM treatment (Fig. 8F b: VII-IX). Further, it was found that different cytokines and chemokines like IL-12 (p40), MCP-1, IL-6, IP-10, KC, RANTES, GM-CSF, MIP-1β, and TNF-α which are known to be associated with CHIKV infection(34) were reduced significantly in the mice serum by TM (Fig. 8I, J **and** K). To further confirm the *in vivo* efficacy of TM on CHIKV infection, survival curve was analyzed. All the CHIKV infected mice died on the 8^th^ day post infection, whereas TM treatment alleviated symptoms and provided complete protection to all the animals (Fig. 8G and H). These results indicate that TM protects mice against CHIKV infection and inflammation.

**Figure 8.**
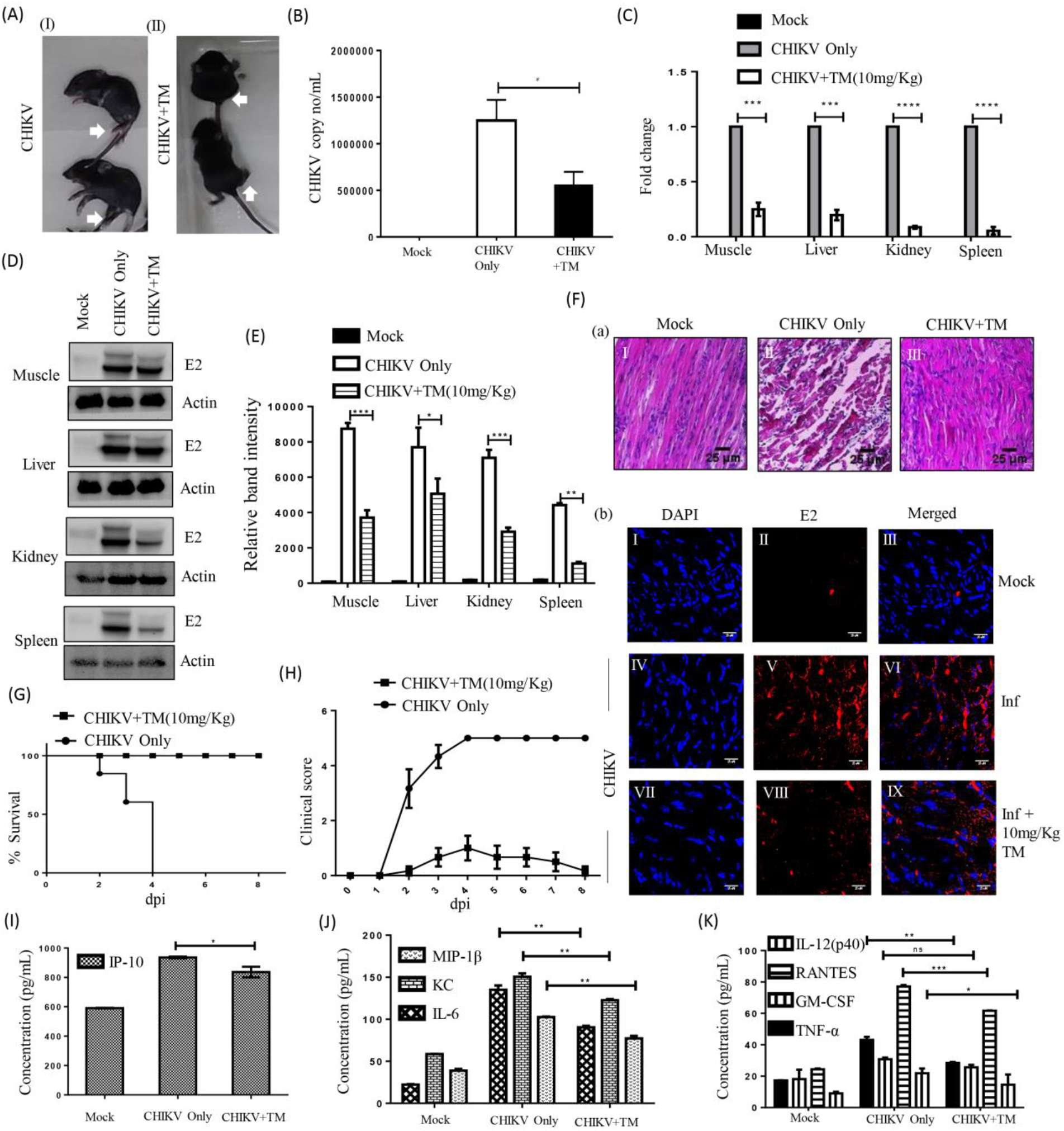
Efficient inhibition of CHIKV infection in mice: C57BL/6 mice were infected subcutaneously with 10^7 pfu of CHIKV-PS and treated with 10mg/Kg TM at every 12h intervals up to 4 or 5dpi. Mice were sacrificed at 5dpi, serum and different tissues were also collected for further downstream experiments. Equal volume of sera were taken to isolate viral RNA. cDNA was synthesized from equal volume of viral RNA and the E1 gene was amplified by qRT-PCR. CHIKV copy number was obtained from standard curve of Ct value. (A) Image of CHIKV-infected as well as drug treated mice. (B) Bar diagram showing CHIKV copy number/mL in virus infected and drug treated mice serum. (C) Whole RNA was isolated from the CHIKV infected and drug treated samples and CHIKV E1 gene was amplified by qRT-PCR. Bar diagram showing the fold change of viral RNA in infected and drug treated samples. (D) Western blot showing the viral E2 protein in different tissue samples. Actin was used as a loading control. (E) Bar diagram showing the relative band intensities of E2 in infected and drug treated samples of different tissues (F) (a) Image panels showing the H&E stained muscles (b) Image panels showing the CHIKV-E2 stained muscles. (G) Survival curve showing the efficacy of TM against CHIKV infected C57BL/6 mice (n=6/group). (H) Line diagram showing the disease symptoms of CHIKV infection which was monitored from 1dpi to 8dpi. (I, J and K) Bar diagrams indicating the concentrations (pg/mL) of different cytokines (IP-10, MIP1-β, KC, IL-6, IL-12(p40), RANTES, GM-CSF, TNF-α) in mock, infected and drug treated mice serum. All bar diagrams were obtained through the Graph pad prism software (n=3 or n=6) Data presented as mean ± SEM. (p ≤ 0.05 was considered statistically significant).

### TM reduces CHIKV infection in hPBMC derived monocyte-macrophage populations *in vitro*

Immunological characterization of hPBMC derived adherent cell population was carried out by staining for specific markers of B cells (CD19), T cells (CD3), monocyte-macrophage cells (CD11b and CD14) and analyzed by flow cytometry (Fig. 9A). It was observed that the adherent population was highly enriched with CD14+CD11b+ monocyte-macrophage cells (Fig. 9A) and was non-toxic up to 100μM concentration of TM (Fig. 9B). The hPBMC derived monocyte-macrophage cells infected with CHIKV and treated with TM (100μM) showed lesser CPE (Fig. 9C). Then, the effect of TM was analyzed using the hPBMC derived monocyte-macrophage cells collected from 3 healthy donors upon CHIKV infection. It was observed that the viral particle formation was reduced by 45% in CHIKV infected hPBMCs in presence of TM (Fig. 9D). The CHIKV infected hPBMCs showed 21.13 ± 1.66% positivity for E2, whereas, pre-incubation with TM for 3h led to 17.4 ± 0.7 % decrease in E2 positive cells (Fig. 9E and F). Taken together, the data suggests that TM might abrogate CHIKV infection significantly in hPBMC derived monocyte-macrophage populations *in vitro*.

**Figure 9.**
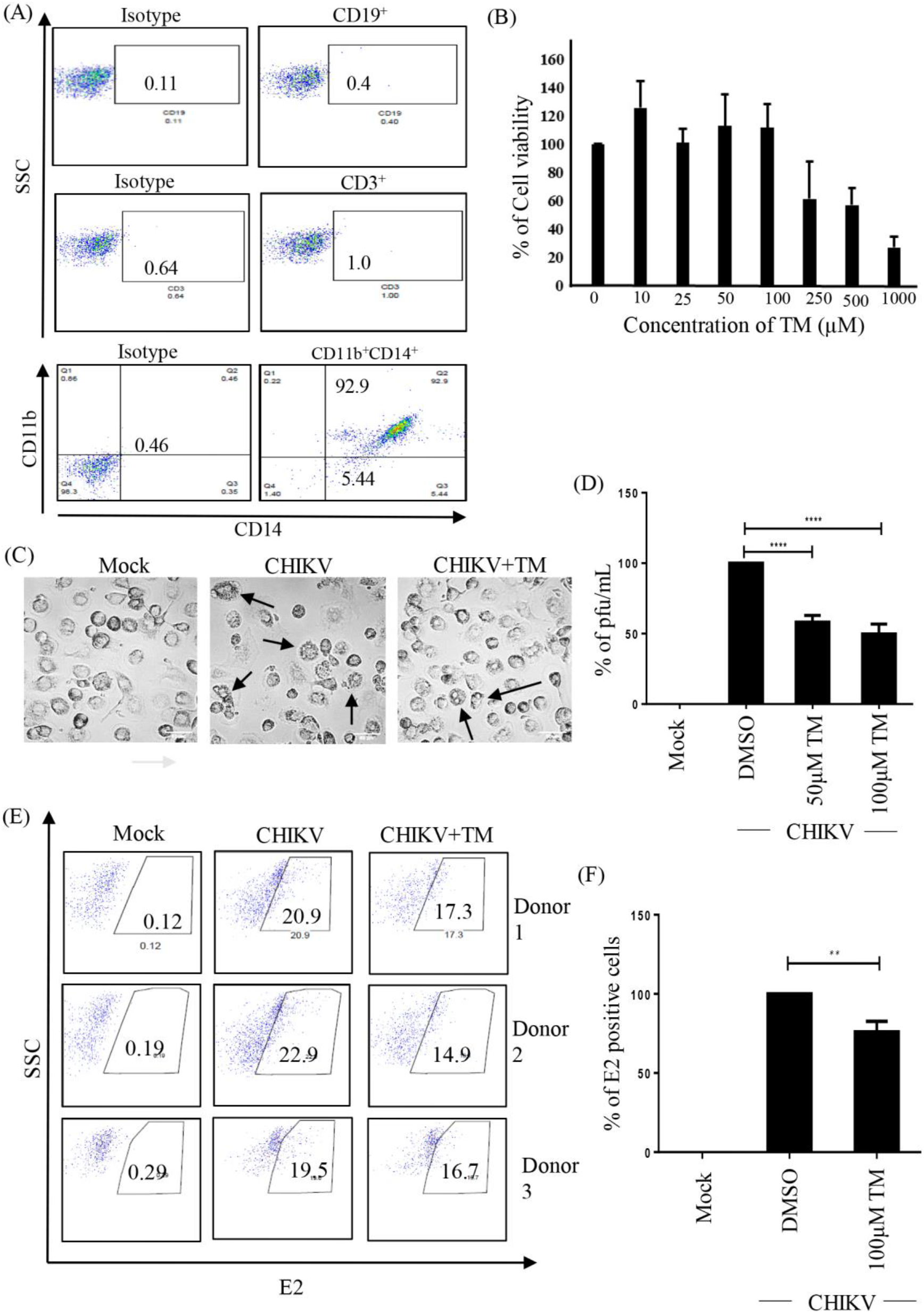
(TM) reduces CHIKV infection in human PBMC (hPBMC) derived monocyte-macrophage populations in vitro. (A) Dot blot showing the percentages of B cells (CD19), T cells (CD3) and CD14+CD11b+ monocyte-macrophage lineage of adherent hPBMCs by flow cytometry. (B) Bar diagram referring the cytotoxicity of TM in hPBMC derived adherent population by MTT assay. (C) Image showing CHIKV induced CPE in hPBMC derived adherent cells. (D) Bar diagram depicting percentage of the viral particle formation by plaque assay. (F) Bar diagram showing the percentage of positive cells for CHIKV E2 protein, determined by Flow Cytometry of CHIKV infected hPBMC derived adherent cells. Data shown are represented as mean ± SEM of three independent experiments. (*, p < 0.05).

## DISCUSSION AND CONCLUSION

CHIKV has now emerged to be an infectious disease of global concern. Unfortunately, there is no effective antiviral for CHIKV. This has pushed efforts for repurposing the existing drugs against CHIKV. Since RAS and PPAR-γ pathways are involved in the progress of viral infections TM, was investigated against CHIKV both *in vitro* and *in vivo*.

It abrogated CHIKV infection efficiently both in the Vero and RAW 264.7 cells with remarkable inhibition in the Viral RNA and protein levels. The reduction in protein level is partly attributed to modulation of the translational control pathway. Additionally, TM interfered in the early and late stages of CHIKV life cycle and was found to be effective in pre and post-treatment assay. Moreover, the agonist of AT1 receptor and antagonist of PPAR-γ increased CHIKV infection that was antagonized by TM. Thus, anti-viral effect of TM could be partly mediated through these host factors. It also diminished the CHIKV induced inflammatory responses. The reduced activation of all the major MAPKs and cytokines by TM through MAPKs inflammatory axis supports the fact that the anti-CHIKV efficacy of TM is partly mediated through the AT1/ PPAR-γ/MAPKs pathways. Further, at the human equivalent dose, TM abrogated CHIKV infection and inflammation *in vivo* leading to reduced clinical score and complete survival of C57BL/6 mice (Fig. 10). Additionally, TM reduced infection in hPBMC derived monocyte-macrophage populations *in vitro*.

**Figure 10.**
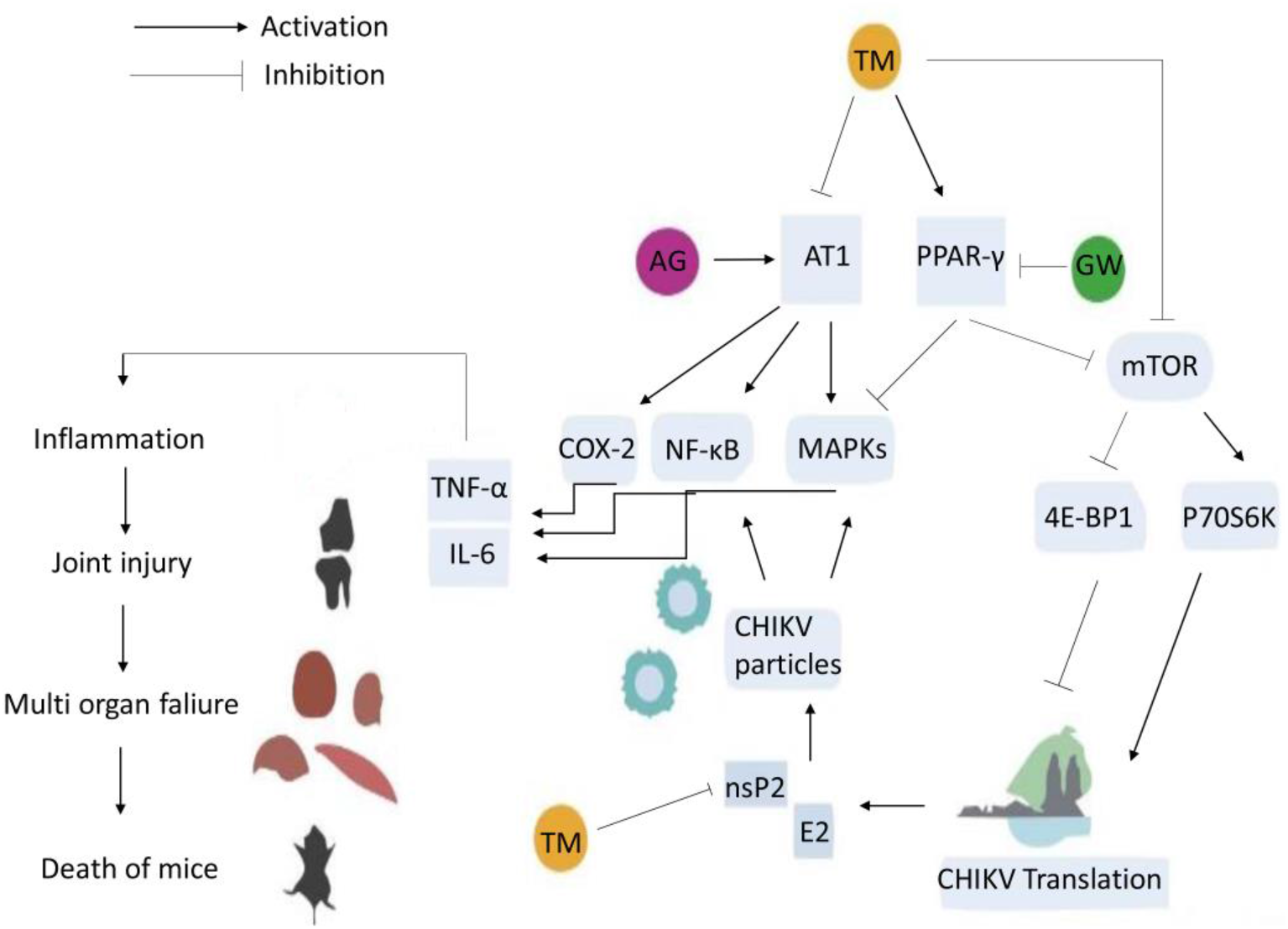
Schematic representation of proposed working model demonstrating the involvement of AT1 PPAR-γ, MAPKs after TM treatment during CHIKV infection. Activation of AT1 and blocking of PPAR-γ induce higher phosphorylation of MAPKs with increase in the expression of COX-2 and NF-κB that lead to increase in the level of inflammatory mediators (TNFα and IL-6). Infection by CHIKV induces these axes to cause inflammation, tissue injury and cell death. Both AG and GW increase CHIKV infection and inflammation. TM blocks AT1 and activates PPAR-γ to reduce CHIKV infection and inflammation. It also modulates the mTOR signaling to interfere in CHIKV translation process leading to significant reduction in viral proteins (E2 and nsP2) and infection as elaborated in the text.

Our preliminary findings have suggested the CHIKV inhibition potential of TM. Subsequently, Tripathi *et.al* have shown inhibition of CHIKV-nsP2 protein by TM with reduction in CHIKV infection. The present study shows its ability to inhibit CHIKV infection by reducing viral RNAs as well as proteins. It is well known that the mammalian target of rapamycin (mTOR) signaling pathway controls protein synthesis in host (39) and is indispensable to viral translation (43). Although the mTOR modulation during CHIKV infection is not well characterized, we have earlier shown increase in expression of mTOR signaling pathway proteins during CHIKV infection *in vitro* (44). TM has been shown to suppress mTOR pathway for its anticancer effect (45). However, similar implication in CHIKV infection is not yet known. Accordingly, by demonstrating remarkable inhibition in the expression of p-mTOR, p-P70S6 kinase and p-4E-BP1 by TM, the present study reveals that this anti-CHIKV property is partly due to modulation of CHIKV protein translation (Fig. 10). Further, this might have impact on the development of arthritis which is often induced by CHIKV infection. This is because arthritis is related to increased mTOR signaling (46) which in turn is associated with reduced PPAR-γ activity (47). As TM is well known as a PPAR-γ activator, the suppression of mTOR signaling might further benefit in alleviation of symptoms of CHIKV.

Moreover, the “time-of-addition” experiment also revealed that TM might interfere in both the early and late stages of the CHIKV life cycle. When treated during infection, there is no effect on the viral particle formation. In contrast, pre and post treatment has greater effect than post treatment only. Thus, TM might not have much effect on viral attachment process and it can be speculated to target multiple phases of CHIKV life cycle to modulate infection. However, this needs further investigation.

Considering the fact that TM is an established AT1 receptor blocker and partial agonist of PPAR-γ, modulation of these host factors might have role in its antiviral efficacy. Thus, AT1 receptor agonist (AG) and PPAR-γ antagonist (GW) were used to demonstrate the involvement of these host factors in the anti-CHIKV property of TM. Evidently, the CPE and levels of viral antigen E2 were high in cells treated with both GW and AG compared to the infection control. This was supported by increase in the CHIKV progeny and structural (E2) as well as non-structural (nsP2) protein levels. Compared to this, treatment of TM significantly reduced CHIKV infection and level of viral proteins. This reveals that activation of AT1 and antagonism of PPAR-γ might be essential for efficient CHIKV infection. This was also evident from up-regulation of AT1 protein level and modest down-regulation of PPAR-γ in infected cells. Further, inhibition of CHIKV infection by PPAR-γ agonist (Rosi) as well as increase in CHIKV infection by PPAR-γ inhibitor (T007) supported the PPAR-γ mediated anti-CHIKV effect of TM (Fig. 10).

In agreement with these actions of TM, the downstream mediators of these axes were also found to be modulated. MAPKs are activated in CHIKV infection (48) and their inhibition has been shown to abrogate its infection (32). Accordingly, the phosphorylation of major MAPKs including p38, ERK1/2 and JNK were observed to be more following infection. While AG and GW induced it further, TM reduced these activated MAPKs significantly. This corroborates that TM partly targets the AT1/ PPAR-γ axes to reduce CHIKV infection. Further, these axes and downstream mediators including MAPKs are also responsible for induction of chemokines and cytokines that are aggravating factors for arthritis. As a result, TM has been shown to reduce inflammation by regulating these mediators (49–52). In the present study, induction of p-NF-κB and COX-2 were found to be significantly reduced by TM, whereas both AG and GW enhanced their expressions. These inflammatory mediators are also demonstrated to be involved in CHIKV induced inflammation with elevated levels of IL-6 and TNF-α in CHIKV-infected macrophage cell lines (28). The levels of the cytokines is often associated with severity of the viral infection (53). Accordingly, treatment with AG and GW increased the levels of TNF-α and IL-6 which was reduced remarkably after the treatment of TM. Since these axes are also mediators of inflammatory process, TM can be expected to manage both infection and symptoms related to CHIKV (Fig. 10).

The *in vitro* findings were validated in C57BL/6 mice model. While infected mice showed symptoms of arthritis and immobility, TM (10mg/Kg, human equivalent dose of 40mg) alleviated these symptoms with reduction in the CHIKV copy number. This was supported by reduction of E1 RNA and E2 protein in different tissues including muscle, liver, kidney and spleen. The reduction in arthritic symptoms, viral copy number, RNA and protein was well supported by the survival curve. The *in vivo* efficacy of TM against CHIKV infection at an acceptable human equivalent dose suggests its suitability for repurposing against this disease. However, it is difficult to extrapolate the pre-clinical data for clinical application without further clinical validation. Nonetheless, to evaluate its suitability, TM was investigated against CHIKV infection in hPBMC derived monocyte-macrophage population *in vitro*. While CHIKV infection increased CPE and E2 positive cells, treatment with TM decreased these effects proving its clinical safety.

The *in vitro* findings, supported by the preclinical data demonstrated the worth of TM to be considered for further investigation for repurposing against CHIKV. Considering the fact that the mTOR as well as AT1/PPAR-γ/MAPK pathways and downstream mediators are modulated by TM, it might regulate both infection and symptoms of CHIKV (Fig. 10). Since these are host pathways, and inflammation is a common etiology in the viral disease progression, TM might have effects against other viruses. In support of this, we found an IC50 of 9.78µM and 26.3 µM against HSV-1 and JEV respectively (Data not shown). This is interesting, since it also suggests that it has a broad spectrum of antiviral action. Also, unlike other AT1 blocker, TM has a better brain access, which makes it suitable for application against viruses affecting brain e.g. CHIKV and JEV. However, it is desirable to reduce the dose of TM by enhancing its bioavailability. Owing to its poor solubility [Biopharmaceutics Classification System (BCS)-II], this may be a challenge (54). Nevertheless, increasing solubility through amorphisation, co-crystallisation or other formulation techniques (55) may enhance its bioavailability and permit a lower dose for application as antiviral.

In conclusion, the antiviral efficacy of TM can be due to its direct and indirect effect on CHIKV targets and host factors. Considering its history of long-term safety and *in vivo* efficacy against CHIKV, it can be repurposed against CHIKV infection after clinical validation.

## ABBREVIATIONS

AT1 receptor: Angiotensin II type 1 receptor
Ang-II: Angiotensin II
ACE: Angiotensin-converting enzyme
AG: [Val5]-Angiotensin II acetate salt hydrate
AF: Alexa Fluor
BSA: Bovine serum albumin
CHIKV: Chikungunya virus
CNS: Central nervous system
CPE: Cytopathic effect
CST: Cell Signalling Technology
COX-2: Cyclooxygenase-2
CPCSEA: The Committee for the Purpose of Control and Supervision of Experiments on Animals
DENV: Dengue virus
DMEM: Dulbecco’s modified Eagle’s medium
DAPI: 4, 6-diamidino-2-phenylindole
dpi: day post infection
EMC: Encephalomyocarditis
ERK: Extracellular signal-regulated protein kinase
ELISA: enzyme-linked immunosorbent assay
4E-BP: eukaryotic translation initiation factor 4E-binding protein
FBS: Fetal Bovine Serum
FACS: Fluorescence-activated cell sorting
GW: GW9662
GM-CSF: Granulocyte-macrophage colony-stimulating factor
hPBMC: human peripheral blood mononuclear cells
HSV: Herpes simplex virus
HRP: Horseradish peroxidase
H & E: Hematoxylin and Eosin
hpi: hour post infection
IL: Interleukin
IS: Indian strain
IRF: Interferon regulatory transcription factor
ICS: Intra cellular staining
IP-10: Interferon gamma-induced protein
10 JEV: Japanese encephalitis virus
JNK: c-Jun N-terminal kinase
KC: Keratinocyte chemoattractant
LASAF: Leica Application Suite Advanced Fluorescence
MAPK: Mitogen-activated protein kinase
mTOR: mammalian target of rapamycin
MTT: 3-(4,5-dimethylthiazol-2-yl)-2,5-diphenyl tetrazolium bromide
MOI: Multiplicity of infection
mAb: Monoclonal Antibody
MCP 1: Monocyte chemoattractant protein 1
MIP-1b: Macrophage inflammatory protein 1-beta
NF-κB: Nuclear factor kappa-light-chain-enhancer of activated B cells
nsP: non-structural protein
NaN_3_: Sodium azide
PPAR-γ: Peroxisome proliferator-activated receptor gamma
PS: Prototype strain
pfu: plaque-forming unit
PBS: Phosphate-buffered saline
pAb: Polyclonal Antibody
PVDF: Polyvinylidene fluoride
PCR: Polymerase chain reaction
q-RT PCR: Quantitative Real-Time Reverse Transcription PCR
RAS: Renin-angiotensin system
RNA: Ribonucleic acid
RPMI: Roswell Park Memorial Institute Medium
RT: Room Temperature
Rosi: Rosiglitazone
RANTES: Regulated upon Activation, Normal T cell Expressed, and Secreted
S6K: ribosomal protein S6 kinase
SDS: Sodium dodecyl sulfate
TM: Telmisartan
T007: T0070907
TNF-α: Tumour Necrosis Factor alpha
WHO: World Health Organization

## ACKNOWLEDGEMENTS

This work was partly funded by Department of Biotechnology (DBT), Ministry of Science and Technology, Govt. of India vide grant no. BT/PR27893/MED/29/1300/2018 and supported by Institute of Life Sciences (ILS), Bhubaneswar core fund provided by DBT, partly by National Institute of Science Education and Research (NISER), Bhubaneswar, Department of Atomic Energy (DAE), Govt. of India and partly by School of Pharmaceutical Sciences, Siksha O Anusandhan Deemed to be University, Bhubaneswar. SD was supported by fellowship from Council of Scientific and Industrial Research (CSIR), Ministry of Science and Technology, Govt. of India. The funding agencies did not have any role in the design of the study and collection, analysis, and interpretation of data and in writing the manuscript.

## CONFLICT OF INTEREST

The authors declare that they have no conflicts of interest.

## DECLARATION OF TRANSPARENCY

This paper adheres to the principles for transparent reporting and scientific rigour of preclinical research as stated in the guidelines for the Design and Analysis, Immunoblotting and Immunochemistry and Animal Experimentation and as recommended by funding agencies, publishers and other organisations involved in supporting research.

## AUTHOR CONTRIBUTIONS

SD, PM, SuC, SoC and BBS conceived the idea, designed the experiments and analyzed the results. SuC, SoC and BBS contributed reagents. SD, SG, SSK, CM, SSP, PM, EL, AR, SC, TM and SK carried out the experiments. SD, PM, SG, SSK, AR, EL, SS, SuC, SoC, BBS wrote & edited the manuscript. SuC, SoC and BBS reviewed the manuscript.

---

**Figure S1.**
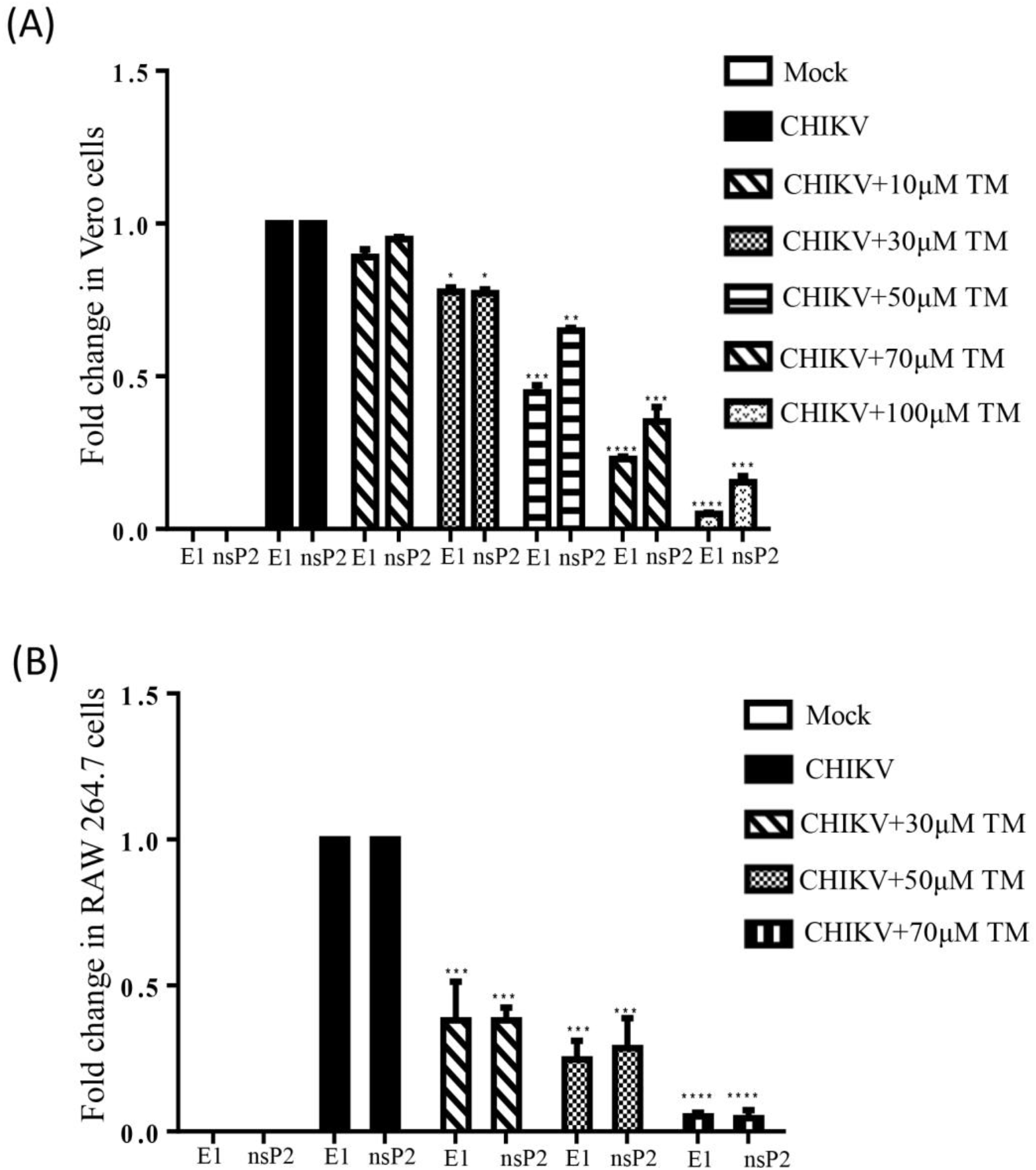
Reduction of CHIKV RNA after TM treatment: Vero cells infected with CHIKV-PS (MOI-0.1) and RAW 264.7 cells infected with CHIKV-IS (MOI-5) were treated with different concentrations of TM and harvested at 18hpi and 8hpi respectively. Whole cell RNA was extracted by TRIzol® and qRT-PCR was performed. (A, B) Bar diagram representing the fold change of E1 and nsP2 genes in CHIKV infected and drug treated Vero and RAW 264.7 cells respectively. Data represented as mean ± SEM (n=3, p ≤ 0.05 was considered statistically significant).

**Figure S2.**
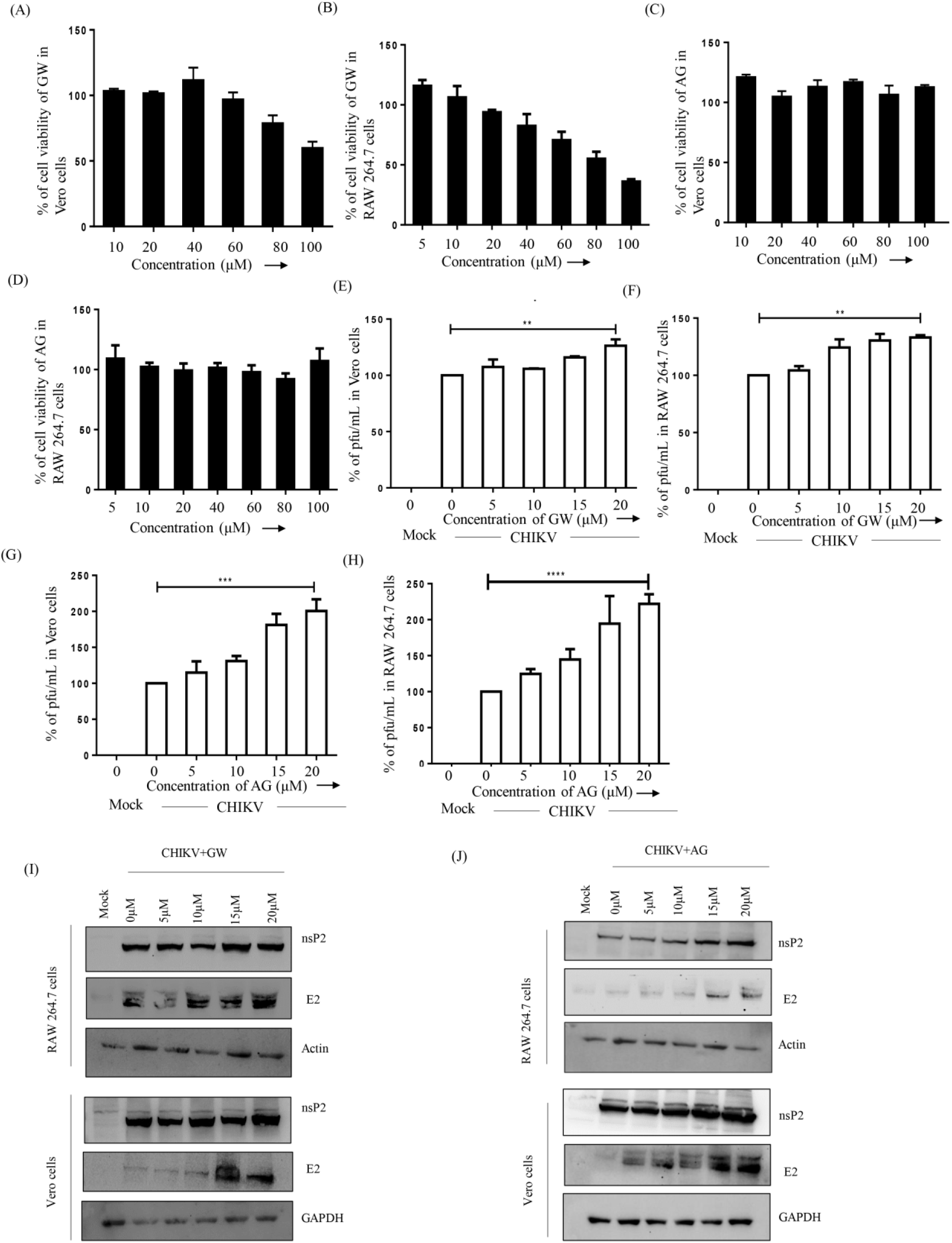
The antagonist of PPAR-γ (GW) and agonist of AT1 (AG) augments CHIKV particle formation dose dependent manner. (A, B) Bar diagram shows the viability of Vero and RAW 264.7 in presence of different concentrations of GW. (C, D) Bar diagram showing the viability of Vero and RAW 264.7 cells in presence of different concentrations of AG. Vero cells were infected with CHIKV-PS and RAW 264.7 cells were infected by CHIKV-IS. After infection GW and AG were added with different concentrations (5µM, 10µM, 15µM, 20µM). The supernatants were collected and virus titers were determined by plaque assay. Cell lysates of the same samples were used for Western Blot. (E, F) Bar diagrams showing the percentage of the viral titer in Vero and RAW 264.7 cells in presence of different concentrations of GW. (G, H) Bar diagrams showing the percentages of the viral titer in Vero and RAW 264.7 cells in presence of different concentrations of AG. Data presented as mean ± SEM. (n=3, p ≤ 0.05 was considered statistically significant). (I, J) Western Blot showing the expressions patterns of CHIKV non-structural and structural proteins.

**Figure S3.**
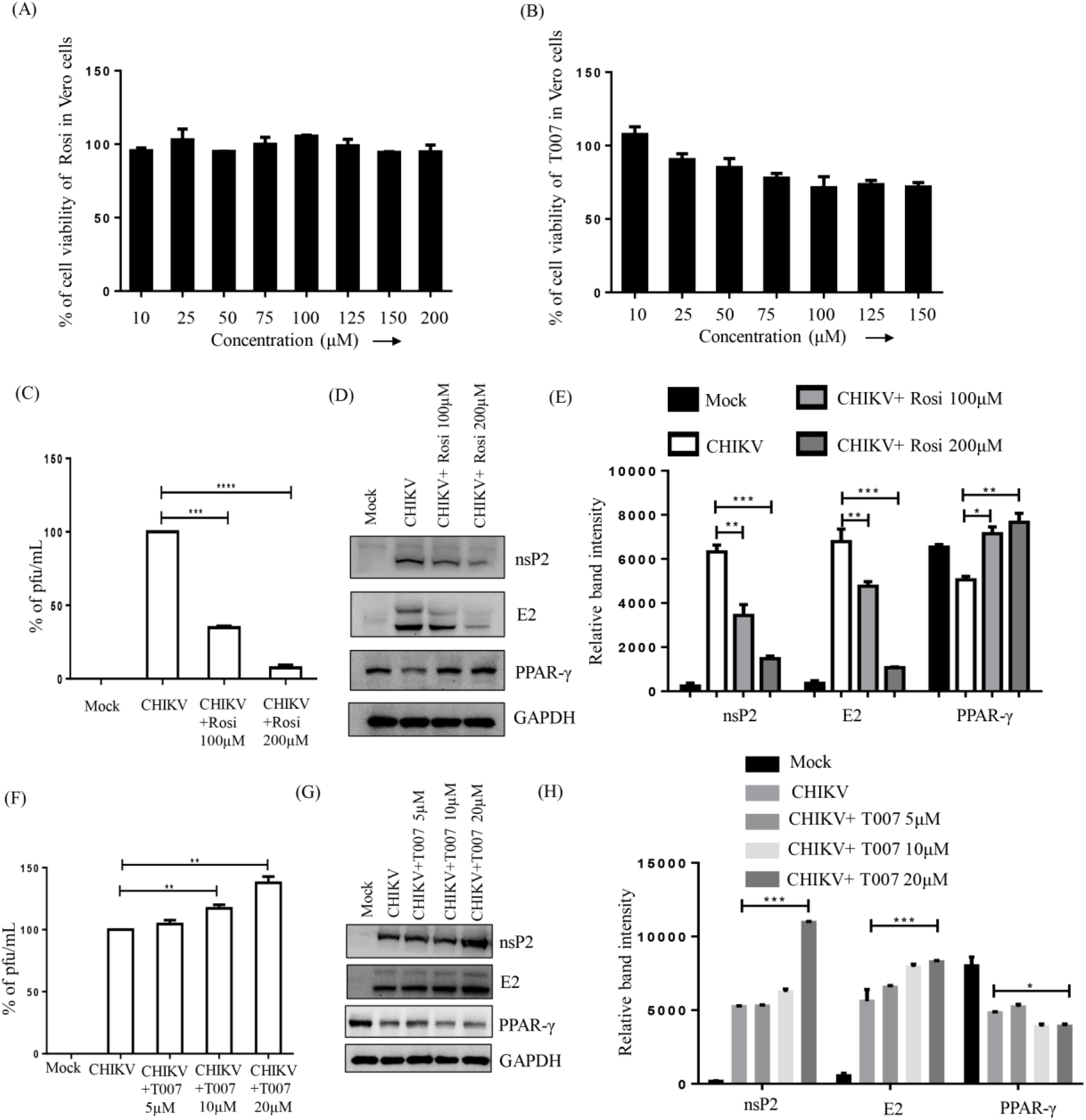
Rosiglitazone (Rosi), the agonist of PPAR-γ, inhibits and T0070907 (T007) the antagonist of PPAR-γ, augments CHIKV particle formation. (A, B) Bar diagrams showing the viability of Vero cells in presence of Rosi and T007. Vero cells were infected by CHIKV-PS at 0.1 MOI and treated with 100 and 200μM of Rosiglitazone (Rosi). Cell supernatants were isolated for plaque assay and Western blot was performed by using the cell lysate. (C) Bar diagrams indicating the percentage of the viral titer in presence of Rosi. (D) Western blot showing the level of PPAR-γ along with viral nsP2 and E2 proteins after Rosi treatment, where GAPDH was used as loading control. (E) Bar diagram depicting the quantitation of nsP2, E2 and PPAR-γ levels. CHIKV-IS infected Vero cells were treated with 50μM T007 and harvested at 18hpi. Plaque assay was performed by using cell supernatant and Western blot by cell lysate. (F) Bar diagram indicating the percentage of the viral titer in presence of T007 (G) Western blot showing the levels of PPAR-γ along with viral nsP2 and E2 proteins after T007 treatment (5μM, 10μM, 20μM), where GAPDH was used as loading control. (H) Bar diagram depicting the quantitation of nsP2, E2 and PPAR-γ levels after different concentrations of T007 treatment. Data presented as mean ± SEM. (n=3, p ≤ 0.05 was considered statistically significant).

**Movie S1: Phenotypic effect of TM in CHIKV infected mice.** C57BL/6 mice were (n=3 in each group) infected subcutaneously with 10^7 pfu of CHIKV-PS and treated with 10mg/Kg TM at every 12h intervals up to 5dpi. Video was taken on 6dpi before sacrifice. The infected group of mice showed symptoms of arthritis in limbs and impaired mobility, whereas TM-treated animals showed no such abnormalities and were moving fine.

**Table S1:**
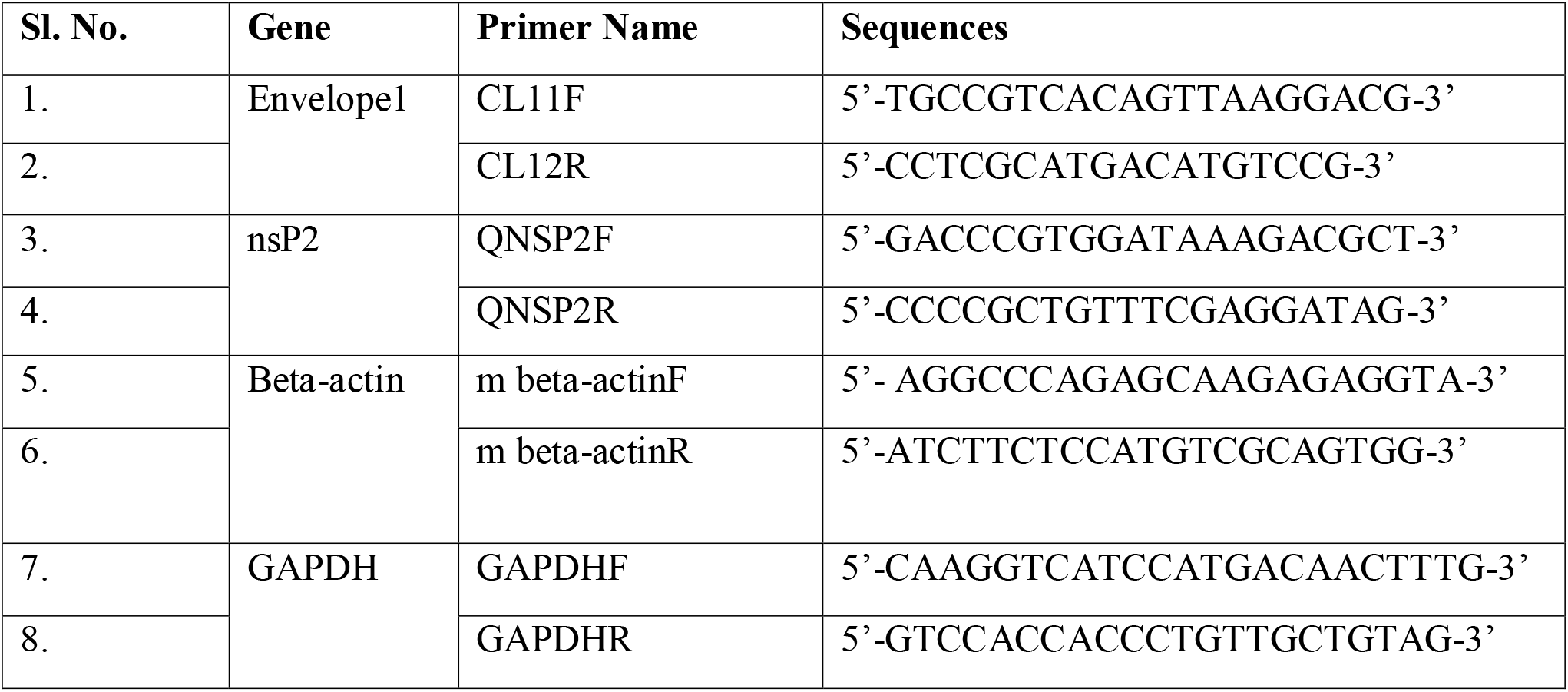
Primer names and sequences.

## Notes

**Data availability statement**: The data that support the findings of this study are available from the corresponding author upon reasonable request. Some data may not be made available because of privacy or ethical restrictions.

